# Astrocytic Cholesterol Fine-Tunes the Balance of Different Modes of Synaptic Exo- and Endocytosis

**DOI:** 10.64898/2025.12.28.696787

**Authors:** Jongyun Myeong, Vitaly A Klyachko

**Affiliations:** Department of Cell Biology and Physiology, Washington University School of Medicine, St. Louis, MO, 63132, United States

**Author notes:** Corresponding author: Vitaly Klyachko.

## Abstract

Cholesterol is essential for organization of neurotransmitter release machinery, yet how it regulates the balance among different forms of synaptic exo- and endocytosis remains poorly understood. Moreover, which pre-synaptic processes rely on neuronal vs astrocyte-derived cholesterol is unknown. Using nanoscale-precision imaging of single-vesicle release in hippocampal synapses we demonstrate that astrocytic cholesterol is a critical determinant of both temporal and spatial aspects of presynaptic dynamics by differentially modulating the two main forms of synchronous release, uni-vesicular (UVR) and multi-vesicular (MVR), effectively fine-tuning their balance. Disruption of astrocytic cholesterol trafficking to neurons combined with its re-supplementation demonstrated that astrocyte-derived cholesterol is necessary and sufficient to determine the UVR/MVR balance. Moreover, astrocytic cholesterol determines the spatial distribution of vesicle release by modulating utilization of different release sites across the active zone. Astrocytic cholesterol also regulates the balance of the two main forms of single-vesicle endocytosis, fast and ultra-fast. These findings suggest that astrocytic cholesterol supply is a critical modulator of synaptic strength that fine-tunes the balance of different forms of synaptic vesicle exo- and endocytosis.

**Highlights:**

- Astrocytic cholesterol determines the balance of uni- and multi-vesicular release
- Cholesterol levels modulate spatial organization of vesicle release
- Astrocytic cholesterol controls the balance of fast and ultra-fast endocytosis
- Astrocytic cholesterol is necessary and sufficient for exo- and endocytosis balance

## INTRODUCTION

Fusion of neurotransmitter-containing vesicles at the synaptic active zone (AZ) critically depends on the composition and dynamics of the presynaptic plasma membrane (PM) (1). Cholesterol is a fundamental component of the PM, modulating the membrane fluidity by inserting its rigid sterol rings between phospholipid molecules, effectively reducing lateral mobility and increasing membrane rigidity (2, 3). The dynamic interplay between cholesterol and phospholipids regulates membrane thickness, compressibility, and permeability to water and ions. At the synapses, cholesterol plays a critical role in the structural organization of PM forming the specialized membrane nanodomains that mediate the clustering of release machinery which orchestrates vesicle release (4-6), including calcium channels and the SNARE complexes. Notably, the supply of cholesterol in the brain is limited to the local biosynthesis by neurons and astrocytes, as cholesterol-delivering lipoproteins from the periphery cannot pass the blood-brain-barrier (7). Adult neurons are believed to be unable to produce sufficient amounts of cholesterol for all their needs and rely on astrocyte-derived cholesterol to maintain some synaptic functions (8, 9). Cholesterol is loaded onto apolipoprotein E (ApoE) in astrocytes and transported by ATP-binding cassette (ABC) transporters A1(ABCA1) and B1 (ABCB1) (10). Secreted lipidated ApoE particles are taken up by neurons via receptor-mediated endocytosis through low-density lipoprotein receptor (LDLR) or LDLR-related protein 1 (LRP1), which are expressed in neuronal PM. Reduced ABC transporter levels or ApoE expression have been implicated in Alzheimer’s disease and other neurological disorders (11), suggesting that astrocytic cholesterol supply is critical for proper neuronal function and survival. However, which synaptic functions and, more specifically, which stages of the neurotransmitter release process require astrocytic cholesterol remains incompletely understood. Moreover, how and what extent spatiotemporal dynamics of synaptic vesicle release depends on astrocytic cholesterol is largely unknown.

Several fundamental forms of vesicle release are present at the synapses, including two major forms of synchronous release known as univesicular (UVR), and multivesicular (MVR), the balance of which determines synaptic strength and is rapidly modulated by neural activity (12, 13). MVR refers to the near-simultaneous release of two or more vesicles, a phenomenon ubiquitously observed in central synapses. This form of release has been suggested to serve a wide range of functions including modulating synaptic reliability/gain, controlling synaptic integration and the induction of several forms of synaptic plasticity (13). We recently developed a nanoscale near-TIRF-based imaging approach that permits analysis of single synaptic vesicle release events in live hippocampal synapses with ∼20 nm precision (14, 15). Using these tools, we found that these two forms of vesicle release have distinctive spatial and temporal organization at the synaptic AZ, in which UVR and MVR utilize partially overlapping but distinct sets of release sites (14). The rapid modulation of these forms of release represents a form of synaptic facilitation widely considered to serve as one of the core mechanisms of information processing (16). Cholesterol is known to be essential for vesicle fusion and cholesterol extraction/depletion has been shown to reduce synchronous release while simultaneously increasing spontaneous release in central neurons (17-19) and the NMJ (20). Yet the role of cholesterol in modulating the organization and balance between the two synchronous forms of neurotransmitter release, and the role of astrocytes in this modulation, have not yet been explored.

Vesicle fusion is rapidly followed by endocytosis and several mechanistically and kinetically distinct forms of endocytosis are present in central synapses (21-25). At the level of individual release events, these include ultra-fast endocytosis (∼50-100 ms) and fast endocytosis (∼0.5-1s), while under high activity conditions, a slow form of endocytosis (tens of seconds) becomes a major contributor to membrane retrieval (24, 26). The utilization of the different forms of vesicle endocytosis varies based on synaptic activity levels and metabolic demands (24). How cholesterol affects different forms of endocytosis is not well understood. Previous studies in the Calyx of Held indicated that kinetics of both fast and slow forms endocytosis are reduced with cholesterol extraction (27), while studies in hippocampal synapses found that spontaneous vesicle endocytosis is increased by cholesterol extraction (17), yet others observed that cholesterol supplementation had no effect on endocytosis (28). Our previous studies have found that single vesicle release events are coupled to either ultra-fast or fast forms of endocytosis in hippocampal synapses, while the slow endocytosis is not detectable (29, 30). How cholesterol in general, and astrocytic cholesterol specifically, modulate the balance and preferential modes of single-vesicle endocytosis has not been examined.

To address these questions, we applied nanoscale measurements of single-vesicle release and uptake in hippocampal synapses to define the role of cholesterol in modulating the balance between different forms of vesicle exo- and endocytosis and the role of astrocytes in this modulation.

## RESULTS

### Cholesterol levels modulate the balance of UVR and MVR

While cholesterol is an important component of the PM regulating membrane fluidity and assembly of exocytotic machinery, how neuronal vs astrocytic cholesterol levels regulate the prevalence and spatiotemporal properties of different forms of synchronous vesicle release (i.e. UVR vs MVR) or vesicle uptake (fast vs ultrafast endocytosis) is unknown. To examine these questions, we used two complementary approaches to alter cholesterol levels either directly at nerve terminals using application of pharmacological agents to acutely extract or supplement neurons with cholesterol (**Figures 1-2**), or indirectly by inhibiting different steps in the pathway of cholesterol transport and delivery from astrocytes to neurons (**Figures 3-4** below).

**Figure 1.**
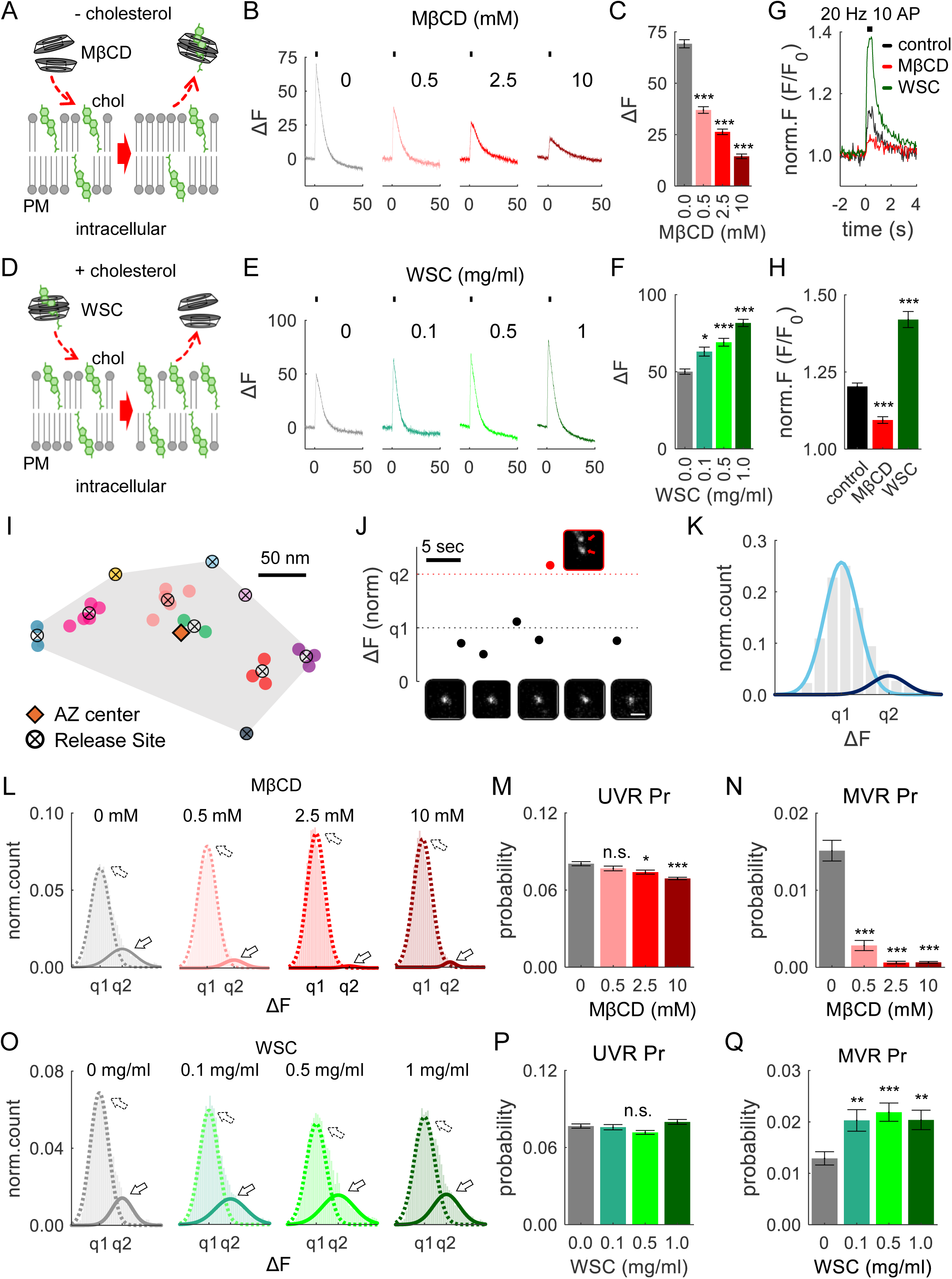
Cholesterol Levels Modulate the Balance of UVR and MVR. **(A)** Schematic of MβCD-mediated cholesterol extraction from the plasma membrane. As MβCD dimer approaches the plasma membrane, cholesterol molecules slide into the hydrophobic cavity of the MβCD dimer, effectively removing cholesterol from the membrane. **(B)** Average vGlut1-pHluorin fluorescence traces during high-frequency stimulation (40 Hz, 50 APs). Neurons were pre-incubated for 10 minutes in solutions containing varying concentrations of MβCD as indicated. **(C)** Amplitude of synaptic responses evoked by high-frequency trains from **(B)** for varying concentrations of MβCD. **(D)** Schematic of cholesterol supplementation using WSC. **(E, F)** Same as (**B, C**), but for different WSC concentrations. **(G, H)** Glutamate release traces in neurons expressing glutamate sensor SF-iGluSnFR(A184S) (G) and quantification of the peak amplitude (**H**) evoked by high-frequency stimulation (20 Hz, 10AP) under cholesterol extraction (10 mM MβCD) or supplementation (1 mg/ml WSC) conditions. **(I)** Sample hippocampal AZ depicting localization of release events evoked by 1 Hz stimulation during 200s. Events occurring at the same cluster/release site are shown in the same color, with cluster centers marked as crossed circles. **(J)** Representative vGlut1-pHluorin fluorescence recording from a single bouton, showing UVR (black) and MVR (red) release events, with corresponding images. Scare bar: 1 µm. **(K)** Amplitude distribution of detected release events revealed two peaks corresponding to q1 (UVR) and q2 (MVR), with Gaussian fit shown in light and dark blue, respectively. **(L)** Amplitude distributions of detected release events with two peaks corresponding to q1 (UVR, dashed line) and q2 (MVR, solid line) at different concentrations of MβCD. **(M, N)** Pr of UVR (**M**) and MVR (**N**) events calculated for individual synapses stimulated at 1 Hz for 200 seconds under varying concentrations of MβCD. **(O-Q)** Same analysis as (**L-N**), but for different concentrations of WSC.

**Figure 2.**
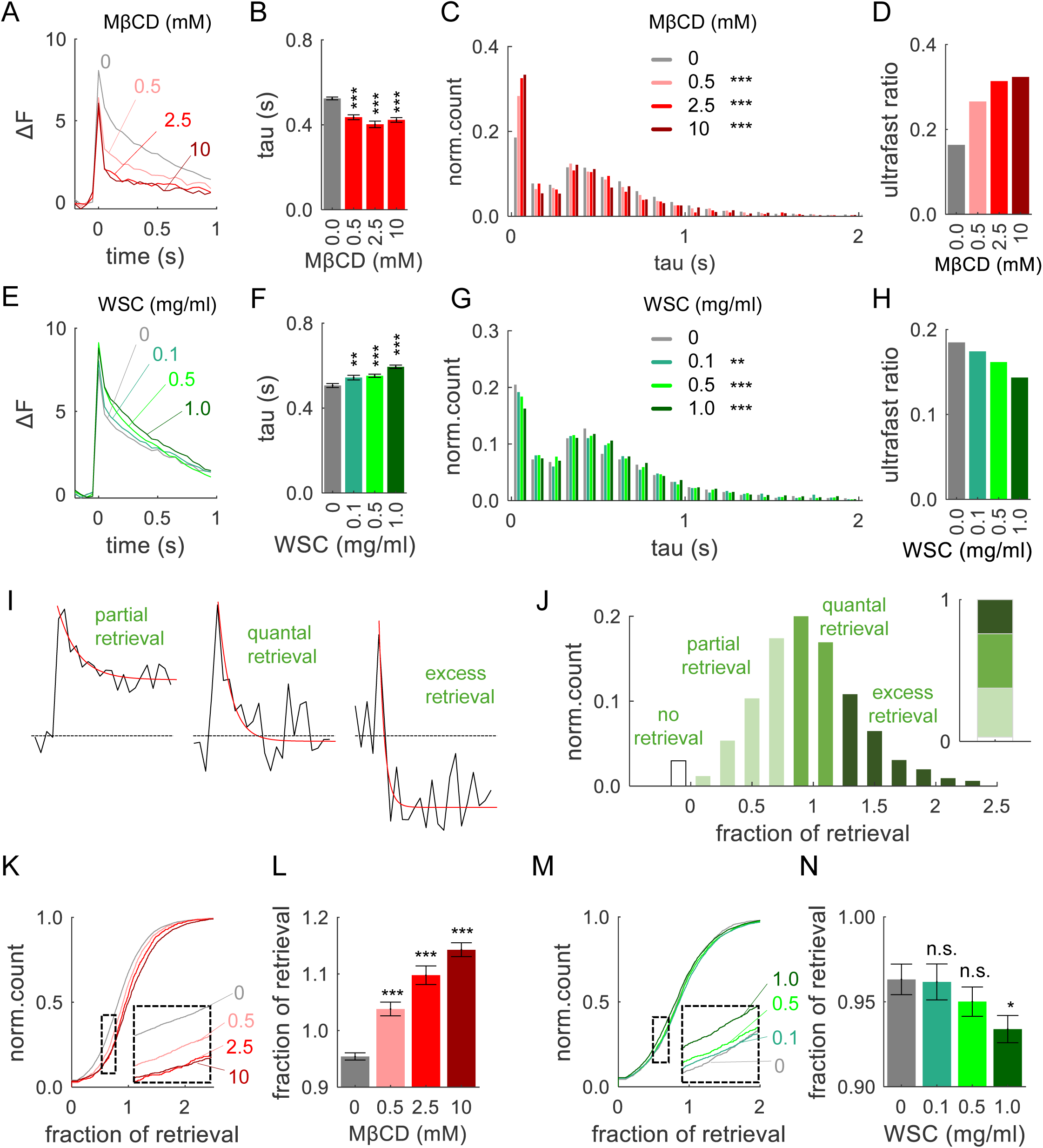
Cholesterol Levels Determine the Balance of Ultra-Fast vs Fast Endocytosis. **(A)** Average single vesicle release traces recorded in different concentrations of MβCD. **(B)** Average release event decay tau in different MβCD concentrations. **(C)** The distribution of individual event decay tau values for 1-AP-evoked release events in different concentrations of MβCD. **(D)** The ratio of ultrafast to fast endocytosis (ultrafast ratio) determined from bi-Gaussian fitting of the distribution of individual event decay tau in (**C**). **(E-H)** Same analysis as (**A-D**), but with varying concentrations of WSC. **(I)** Representative examples of single release events displaying different fractions of retrieval with corresponding fitting traces (red lines). **(J)** Histogram of fractions of retrieval following single-AP stimulation and its quantification into four groups (inset). **(K)** Cumulative plot of retrieval fractions under different concentrations of MβCD. **(L)** Average retrieval fractions under different concentrations of MβCD. **(M, N)** Same analysis as (**K, L**), but with different concentrations of WSC.

**Figure 3.**
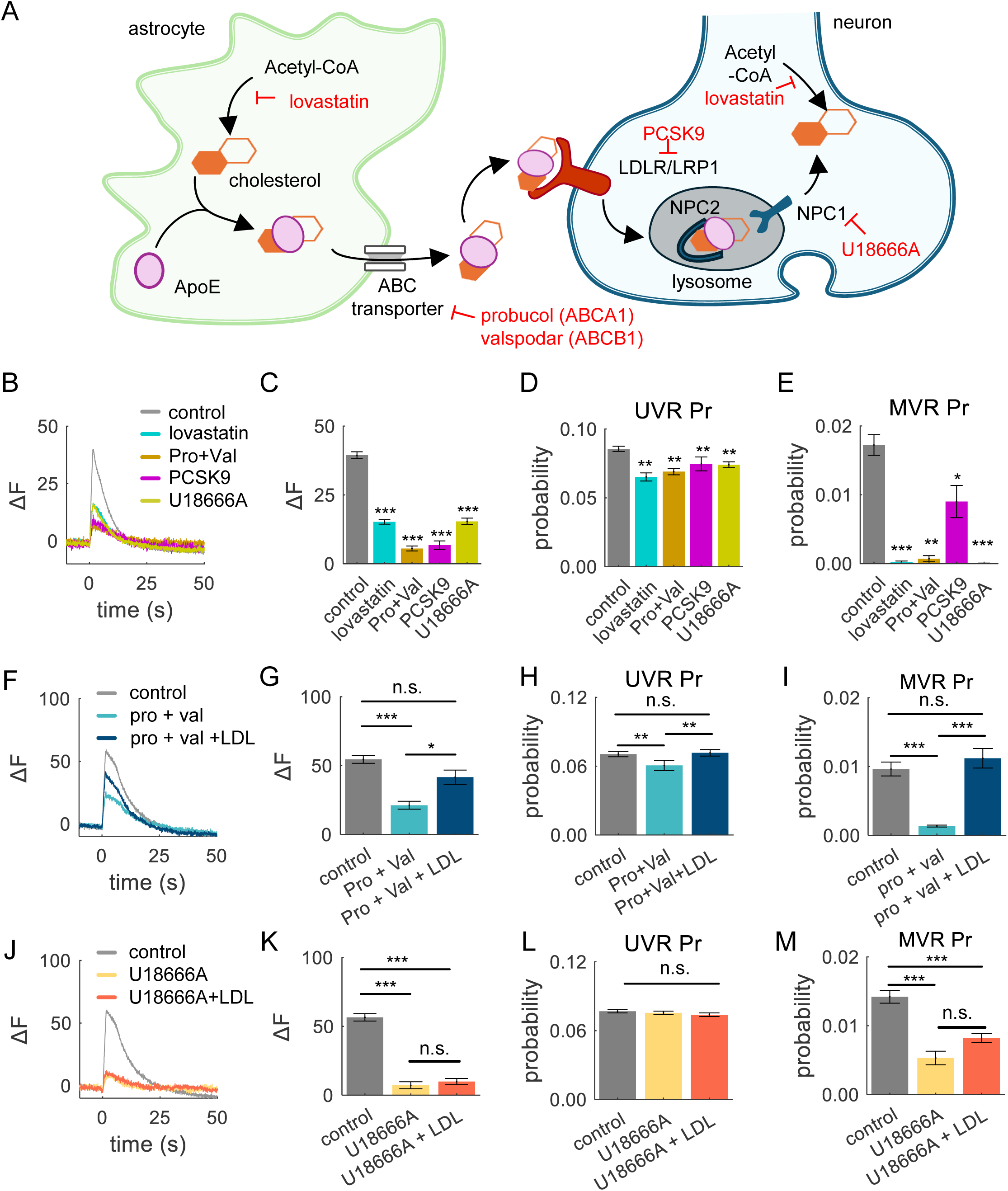
Astrocytic Cholesterol Fine-tunes the Balance of Different Forms of Exocytosis. **(A)** Schematic pathway of cholesterol synthesis, secretion, and transfer from astrocytes to neurons. Cholesterol synthesis was blocked by lovastatin (10 µM), while astrocytic lipoprotein efflux via ABCA1 and ABCB1 was inhibited by probucol (10 µM) and valspodar (5 µM), respectively. The uptake of lipoproteins via LDLR and LRP1 was inhibited by PCSK9 (50 nM). The dissociation of internalized lipoprotein in the lysosomes by NPC1 was inhibited by U18666A (5 µM). **(B)** Average vGlut1-pHluorin fluorescence traces evoked by high-frequency stimulation (50AP at 40Hz) when either cholesterol synthesis, astrocytic secretion, neuronal uptake, or lysosomal dissociation was inhibited for 2 days prior to recording, as indicated. **(C)** Quantification of the peak vGlut1-pHluorin amplitude in data from (**B**). **(D, E)** Pr of UVR (**D**) and MVR (**E**) in individual synapses, evoked at 1 Hz for 200 s under the same conditions as in (**B-C**), as indicated. **(F-I)** Same set of analyses as (**B-E**), but for probucol and valspodar treatment with or without 10 µg/mL of LDL supplementation for 2 days. **(J-L)** Same set of analyses as (**B-E**), but for U18666A treatment with or without 10 µg/mL of LDL supplementation for 2 days.

**Figure 4.**
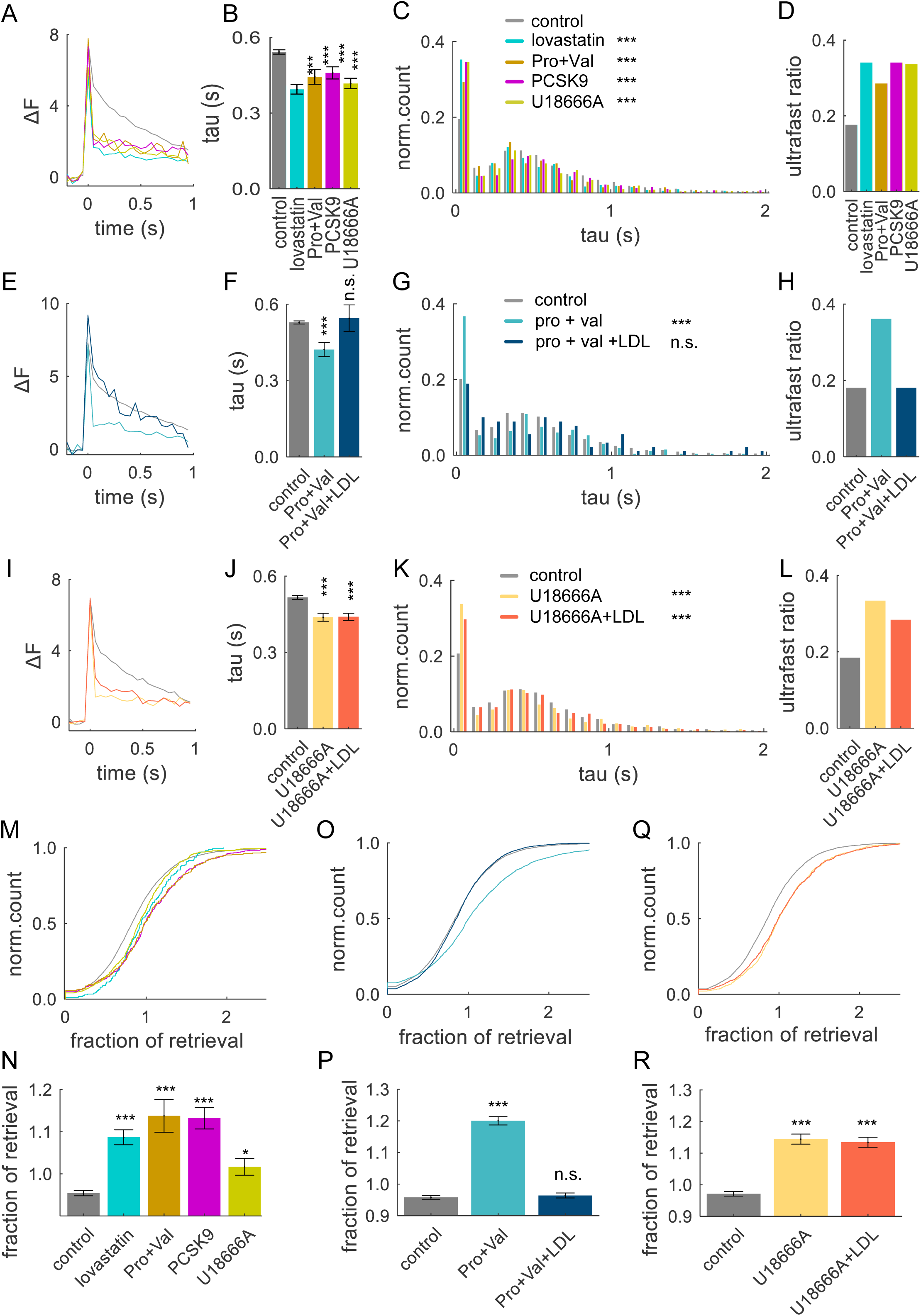
Astrocytic Cholesterol Modulates the Balance of Fast and Ultra-Fast Endocytosis. **(A)** Average single-event traces recorded following a 2-day pre-incubation with various inhibitors that disrupt cholesterol supplementation from astrocytes to neurons as shown in **Fig 3A**. Control - black; lovastatin (10 µM) - cyan, probucol (10 µM) and valspodar (5 µM) – orange; PCSK9 (50 nM) - purple, U18666A (5 µM) – green. **(B)** Average decay kinetics of individual events in (**A**) in different inhibition conditions. **(C)** Distribution of event decay tau values for single release events under the same inhibition conditions as in (**A**). **(D)** The ultrafast/fast endocytosis ratio determined from fitting the event decay distributions in (**C**) in different conditions indicated. **(E-H)** Same analyses as (**A-C**), but with probucol and valspodar co-incubation, with or without 10 µg/mL LDL supplementation, for 2 days. **(I-L)** Same analyses as (**A-C**), but with U18666A pre-incubation, with or without 10 µg/mL LDL supplementation, for 2 days. **(M, N)** Cumulative plots (**M**) and average values (**N**) of fractional retrieval in synapses under different inhibition conditions as indicated. **(O, P)** Same analyses as (**M, N**), but for probucol and valspodar co-incubation with or without 10 µg/mL LDL supplementation, for 2 days. **(Q, R)** Same analyses as (**M, N**), but for U1866A pre-incubation with or without 10 µg/mL LDL supplementation, for 2 days.

In the first set of experiments, to acutely manipulate PM cholesterol levels, we applied variable concentrations of cholesterol-chelating agent methyl-beta-cyclodextrin (MβCD) to reduce cholesterol levels at the synaptic PM, or a preloaded MβCD-cholesterol complex called “water-soluble cholesterol” (WSC) to increase PM cholesterol levels (**Figure 1A and 1D**). To minimize unintended longer-term effects of these alterations on cellular metabolism, such as triggering intracellular cholesterol transport to/from the PM (31, 32), application of both agents was limited to 10 min immediately prior to our measurements. To assess vesicle release, a pH-sensitive indicator pHluorin was targeted to the synaptic vesicle lumen via vGlut1 (vGlut1-pHluorin) (25, 33) using lentiviral infection at DIV3 in cultured hippocampal neurons.

We first determined the effectiveness of altering cholesterol levels by examining synaptic responses elicited in individual synapses by high-frequency stimulation (40 Hz, 50 AP). Increasing MβCD concentration and thus reducing PM cholesterol led to a progressively greater reduction in synaptic response to high-frequency stimulation (**Figure 1B,C**; statistical data for all experiments is provided in **Table S1**), in agreement with the established role of cholesterol in maintaining synaptic vesicle release in neurons (17-19). We observed an opposite effect when cultures were instead supplemented with cholesterol using WSC, which progressively increased synaptic response (**Figure 1E,F**). We considered a possibility that experimentally altering cholesterol levels could affect synaptic response via an effect on vGlut1-pHluorin sensitivity or other properties by altering its interactions with the membranes. We thus sought to validate the observed effects with an independent approach and performed imaging of glutamate release at single synapses using glutamate sensor SF-iGluSnFR(A184S) (34) selectively expressed in neurons. In agreement with the experiments above, acute cholesterol extraction (with 10mM MβCD) strongly reduced glutamate release **(Figure 1G,H)**, whereas cholesterol supplementation (with 1 mg/ml WSC) enhanced glutamate release during 20 Hz 10 AP stimulation.

We then used these approaches to examine how altering cholesterol levels affect the occurrence of different forms of vesicle release evoked by single action potentials (APs). Release events were detected and localized with ∼20 nm precision at the individual AZs, using a combination of near-TIRF imaging with well-established computational detection and localization approaches at 37°C, recorded at 50 ms/frame, as previously described (**Figure 1I**) (15, 30). Low-frequency stimulation at 1 Hz for 200 s evoked both UVR and MVR events, the two principal forms of synchronous release (**Figure 1J**). In our measurements, MVR was commonly detected as the simultaneous release of two vesicles, corresponding to a second peak in the quantal event amplitude distribution (**Figure 1K**). As in experiments above, we altered cholesterol levels using a 10 min pre-incubation with different MβCD or WSC concentrations to progressively reduce or increase synaptic PM cholesterol levels, respectively. We first determined the basal vesicle release probability (Pr) and found that acute reduction of cholesterol levels with increasing MβCD concentrations resulted in progressively reduced Pr (**Figure S1A**). Cholesterol supplementation through WSC had an opposite (and somewhat weaker) effect - increasing the Pr with the increase in WSC concentration (**Figure S1B**), in line with our observations using high-frequency trains (**Figure 1C and 1F**). Next, we asked whether cholesterol differentially modulates the two main forms of synchronous release, UVR and MVR. To assess the relative prevalence of the two forms of release, we plotted event probability distribution and applied a bi-Gaussian fitting approach to quantify the proportion of UVR and MVR events in normal conditions (**Figure 1K**), and in cholesterol-reduced or -increased conditions (**Figure 1L and 1O**, respectively). We observed a strong progressive decrease in the occurrence of MVR with decrease in cholesterol levels by MβCD (**Figure 1L and 1N**), with nearly complete abolishment of MVR (>14 fold reduction) at higher MβCD concentrations (>2.5 mM). In contrast, cholesterol supplementation via WSC resulted in a significantly increased proportion of MVR events (**Figure 1O and 1Q**). Notably, in both conditions, UVR was much less sensitive to changes in cholesterol levels than MVR, with a reduction of only 15% at highest MβCD concentration (1400% for MVR), and no significant increase (4%) at the highest WSC levels (was 36% for MVR) (**Figure 1M and 1P**). Thus, while both forms of synchronous release are regulated by cholesterol levels, the occurrence of MVR is highly sensitive to the cholesterol alterations, while UVR is much less affected. Thus, changes cholesterol levels could be a potent factor modulating the MVR/UVR balance (**Figure S1C,D**).

In addition to the two synchronous forms of release, asynchronous release that persists for tens to hundreds of milliseconds after an AP is also known to regulate several major aspects of synaptic function (12). Cholesterol has been reported to regulate asynchronous release in the crayfish neuromuscular junction (35). We performed detection of asynchronous release events in our measurements, and found that Pr of this form of release is also somewhat sensitive to changes in cholesterol levels in hippocampal neurons, with sensitivity similar to that of UVR (**Figure S1E,F**).

These results suggest that cholesterol levels modulate the balance of the two main components of synchronous release with the occurrence of MVR being several folds more sensitive to cholesterol levels than other forms of release.

### PM Cholesterol Regulates the Spatial Distribution of Vesicle Release Across the AZ

Recent studies have shown that vesicle release is not randomly distributed across the AZ but follows a precise spatiotemporal organization, characterized by the repeated utilization of several specialized release sites (29, 30, 36, 37). We thus next asked whether the spatial organization of vesicle release is dependent on cholesterol levels. To assess spatial properties, we measured distances between each release event and all other events in the same AZ as well as from each release event to the AZ center. The AZ area was defined as the convolute hull encompassing all detected release events at that AZ, and its centroid was used to define the AZ center (**Figure 1I**). This “functional” AZ definition closely matches the ultrastructurally defined AZ dimensions observed by electron microscopy (38). Cholesterol sequestration with MβCD caused release events to occur closer to the AZ center, in a concentration-dependent manner (**Figure S2A,B**) and, accordingly, to occur at shorter distances from each other compared to control conditions (**Figure S2C,D**). In contrast, cholesterol supplementation via WSC at 1mg/mL (but not at lower concentrations) caused the events to occur further away from the AZ center (**Figure S2E,F**), and release events were localized farther apart than in control conditions (**Figure S2G,H**). We further examined how altering cholesterol levels influenced the number of release sites utilized at individual synapses. We applied a previously established hierarchical clustering algorithm with a 50-nm clustering diameter to define individual release event clusters, which may be interpreted to represent release sites within each AZ (**Figure 1I**). Cholesterol sequestration significantly reduced the number of release sites utilized per individual synapse (**Figure S2I)** and the number of events detected per release site also decreased **(Figure S2J**), whereas cholesterol supplementation had an opposite effect (**Figure S2K,L**).

Together these findings suggest that PM cholesterol levels modulate the spatial distribution of vesicle release across the AZ and utilization of vesicle release sites.

### Cholesterol Levels Modulate the Balance of Different Modes of Endocytosis

Vesicle fusion is immediately followed by endocytosis that rapidly retrieves synaptic vesicle components through several distinct pathways. Previous studies have identified several forms of endocytosis coupled with single-vesicle release events, including ultrafast endocytosis with a decay time constant of ∼50-100 ms and fast endocytosis with a time constant of 0.5-1 s (21-23, 29, 33). In our measurements, the kinetics of the pHluorin signal following vesicle fusion primarily reflect the endocytosis process (15, 21, 29, 30, 33, 39). To assess the impact of cholesterol on endocytosis kinetics, we first examined the averaged traces of single vesicle release events after systematically decreasing or increasing PM cholesterol (**Figure 2A and 2E**). We found that MβCD-mediated cholesterol extraction strongly accelerated the average decay kinetics of release events, while cholesterol supplementation had an opposite effect (**Figure 2A,B and 2E,F**). To assess if these changes are caused by a shift in the preferential mode of vesicle retrieval (i.e. ultrafast vs fast modes of endocytosis), we analyzed the distributions of endocytosis kinetics across the release event population. We applied a mono-exponential fit to approximate the decay kinetics for each event trace (**Figure S3A**), revealing that release events had one of the two types of decay kinetics: ∼100 ms (ultrafast endocytosis) or ∼500 ms (fast endocytosis). In agreement with earlier findings (21, 30), we observed two peaks in the distribution of events’ decay tau (**Figure 2C**), with fast endocytosis accounted for ∼80% of events and ultrafast endocytosis for ∼20%. Fitting the two peaks of the Gaussian distribution provided quantification for the proportion of ultrafast-to-fast forms of endocytosis across the event population, which was quantified as an “ultrafast” ratio (**Figure 2D**). We then quantified the effects of altering cholesterol levels on the proportion of ultra-fast vs fast forms of endocytosis. Cholesterol depletion with MβCD caused a marked shift in the event decay distribution towards the ultra-fast component and a corresponding increase in the “ultrafast” ratio (**Figure 2C,D**). Cholesterol supplementation with WSC had an opposite effect with a shift towards more fast endocytosis (**Figure 2G**), accompanied by a reduction in ultrafast ratio (**Figure 2H**).

In addition to the decay kinetics, endocytosis can be characterized by the dwell time (**Figure S3B**), which reflects a variable delay often observed in the initiation of endocytosis process. We observed that reducing PM cholesterol level shifted the distribution of dwell time towards shorter durations and significantly reduced the average dwell time (**Figure S3C**), while cholesterol supplementation had an opposite effect (**Figure S3E**). Similarly, the event half-time, which is a broader measure of event decay duration, was significantly shorter upon cholesterol reduction and longer upon cholesterol supplementation (**Figure S3D,F**).

The extend of endocytosis following single release events varies widely within and across synapses, ranging from partial/incomplete to complete (i.e. quantal) and, in some cases, even excessive (**Figure 2I**). We categorized single-vesicle events into these three categories based on the remaining event amplitude at the end of the 1s interval following and AP, normalized to the immediately preceding baseline. To mitigate the effects of noise in single vesicle recordings, events which decay to 0.8-1.2 of the baseline were classified as complete/quantal, those with a decay <0.8 as incomplete, and those >1.2 as excessive. In our recordings, ∼38% of events exhibited a complete retrieval, while 35% showed partial retrieval, and ∼24% displayed excessive retrieval (**Figure 2J**). The remaining ∼3% did not show measurable retrieval during 1-second interval between stimuli, and were not included in the subsequent analysis. Within these definitions, we quantified the effects of altering cholesterol levels on the extent of vesicle retrieval and found that cholesterol reduction resulted in a significant shift toward more excessive retrieval, which progressively increased with larger MβCD concentrations (**Figure 2K,L and S4A**). In contrast, supplementation of cholesterol with WSC had an opposite effect with a shift towards more partial retrieval in a concentration-dependent manner (**Figure 2M,N and S4B**).

Taken together, these findings indicate that PM cholesterol level is an important determinant of the mode and extent of single-vesicle endocytosis following 1-AP stimulation.

### Astrocytic Cholesterol Modulates the Balance of Different Forms of Exocytosis and the Spatial Distribution of Vesicle Release

Due to the restrictive nature of the blood-brain barrier, which prevents the uptake of circulating cholesterol (7), most of cholesterol in the brain is synthesized locally (8-10). While both neurons and astrocytes contribute to cholesterol synthesis, neurons are believed to be unable to produce sufficient amounts of cholesterol to sustain their needs (40, 41). Thus most of neuronal cholesterol is initially synthesized by astrocytes and is transported to neurons via lipoproteins, primarily in the form of APOE-bound cholesterol (APOE-chol) which is secreted via astrocytic ABCA1 and ABCB1 transporters (**Figure 3A**). APOE-chol is then taken up by neurons via LDLR or LRP1 receptors, and transported to neuronal lysosomes, where it is dissociated by Niemann-Pick disease, type C1 (NPC1) and C2 (NPC2) proteins. How different forms of synaptic vesicle exo- and endocytosis depend on astrocytic cholesterol is poorly understood.

We sought to address this question by targeting cholesterol production or different steps in the pathway of astrocytic cholesterol transport and uptake by neurons. In the first set of experiments, we incubated cultures for 2 days with 10 µM lovastatin to inhibit cholesterol synthesis in both neurons and astrocytes to establish the baseline requirements for the naturally produced cholesterol in synaptic transmission. In subsequent experiments, to define the specific roles of astrocytic cholesterol in the balance of different forms of exo-/endocytosis, we targeted different steps in the astrocytic cholesterol production/secretion/uptake either using 10 µM probucol and 5 µM valspodar to block astrocytic cholesterol ABCA1 and ABCB1 transporters, or using 50 nM recombinant PCSK9 to block LDLR and thus APOE-chol uptake into neurons, or using 5 µM U18666A to inhibit cholesterol dissociation from APOE in neuronal lysosomes. All of these treatments strongly reduced synaptic vesicle release evoked by high-frequency stimulation **(Figure 3B,C)**, as well as the probability of single-vesicle release events (**Figure S5A**). Most importantly, disruption of astrocytic cholesterol by each of these treatments had a several fold larger effect on the probability of MVR than UVR (**Figure 3D,E**), resulting in a markedly reduced MVR/UVR ratio (**Figure S5B**), in a close agreement with our initial cholesterol extraction/supplementation experiments. These results are also consistent with the observations that occurrence of MVR was greatly reduced in neuronal cultures grown in the absence of an astrocyte feeder layer (although some amount of astrocytes was likely still present in the culture) (**Figure S5C-E**). Notably the effects of lovastatin that inhibits cholesterol production in both neurons and astrocytes were comparable in magnitude to the effects of selectively inhibiting astrocytic cholesterol transport or uptake, supporting the notion that synaptic cholesterol originates predominantly from astrocytes. Together these results suggest that astrocytic cholesterol is a major determinant of the MVR/UVR balance.

We next examined if external cholesterol is not only necessary, but also sufficient to maintain synaptic transmission and the UVR/MVR balance. We blocked cholesterol secretion from astrocytes for 2 days by inhibiting ABCA1 and ABCB1 transporters while simultaneously supplying exogenous LDL (10 µg/ml) to determine if it sufficient to rescue synaptic release deficits in the absence of astrocytic cholesterol secretion. Exogenous LDL was able to largely rescue synaptic transmission deficits caused by ABC transporter inhibition during high-frequency stimulation (**Figure 3F,G**). Similarly, the Pr of both UVR and MVR evoked by single AP-stimulation was restored upon LDL supplementation (**Figure 3H,I)** and MVR/UVR balance was also fully normalized **(Figure S5F**). We confirmed that this normalization was mediated by uptake of exogenous cholesterol rather than by stimulating neuronal cholesterol synthesis, because no rescue by LDL was observed when NPC1 was inhibited instead (**Figure 3J-M and S5G**), which is a protein required for the endocytosed cholesterol to be properly processed/exported from the neuronal lysosomes, but not for the neuronal cholesterol production.

In addition to modulating the mode of vesicle release, our results above suggested that changes in cholesterol levels modulate the spatial distribution of vesicle release across the AZ (**Figure S2**). We thus analyzed the spatial localization of release events under conditions when different steps in the astrocytic cholesterol transport/endocytosis/lysosomal export were blocked, to determine if astrocytic cholesterol is necessary and/or sufficient for this modulation. Inhibiting cholesterol synthesis in general, or different steps in astrocytic cholesterol pathway more specifically, caused a significant shift in the release event localization towards the AZ center (**Figure S6A,B**) and a corresponding shortening of the event-to-event distance (**Figure S6C,D**), in a close agreement with the spatial effects of direct cholesterol extraction. As in experiments above, these spatial changes were fully normalized by supplementation of exogenous LDL (**Figure S6E-S6H**), but not under conditions when NPC1 was inhibited instead (**Figure S6I-S6L**), supporting the notion that external/astrocytic cholesterol is necessary and sufficient to set the spatial localization of vesicle release across the AZ.

Taken together, these results suggest a critical role of astrocytic cholesterol in supporting synaptic vesicle release, its spatial organization, and its particularly critical role in fine-tuning the MVR/UVR balance.

### Astrocytic Cholesterol Modulates the Balance of Fast and Ultra-Fast Endocytosis

Finally, we examined the role of astrocytic cholesterol in modulating the balance of different forms of endocytosis. Broad inhibition of cholesterol synthesis in neurons and astrocytes, or all of the above treatments that more specifically target the pathway of astrocytic cholesterol transport/uptake/processing in neurons, resulted in a significantly faster average decay kinetics of individual events (**Figure 4A,B**). All of these treatments also caused a corresponding shift in the distribution of single event decay kinetics towards increased ultrafast endocytosis (**Figure 4C**), and a ∼2-fold increase in ultra-fast ratio (**Figure 4D**). These treatments similarly caused an increased proportion of excessive retrieval (**Figure 4M,N**). The extent of changes in endocytosis caused by inhibition of astrocytic cholesterol were comparable to changes caused by broad inhibition of cholesterol synthesis in both astrocytes and neurons, and to the extent of changes caused by cholesterol extraction. This result supports the notion that astrocytic cholesterol is the predominant source of cholesterol for neurons and is a critical determinant of the balance of different forms of endocytosis. We further established that extracellular/astrocytic cholesterol was sufficient to set the ultra-fast/fast endocytosis ratio since the changes in endocytosis caused by blocking astrocytic ABCA1 and ABCB1 transporters were fully normalized by exogenous LDL supplementation (**Figure 4E-H and 4O,P**), while exogenous LDL failed to normalize changes in endocytosis balance when NPC1 was inhibited instead (**Figure 4I-L and 4Q,R)**. Taken together these results suggest that astrocytic cholesterol determines the balance of different forms of endocytosis.

## DISCUSSION

Cholesterol is widely known to be essential for organization and function of neurotransmitter release machinery at synapses (1, 6). Yet how it regulates different forms of synaptic vesicle exo-and endocytosis and their spatiotemporal organization in small central synapses has remained poorly understood due to lack of direct measurements of these processes at individual synaptic boutons. Moreover, to what extent neuron-derived vs astrocyte-derived cholesterol is required for various vesicle release/uptake mechanisms remains largely unknown. Here we used a nanoscale-precision imaging of individual vesicle release events evoked by single APs in hippocampal synapses to examine the role of cholesterol, and specifically of astrocytic cholesterol, in modulating fundamental spatiotemporal properties of synaptic exo- and endocytosis. Our results indicate that cholesterol supply, and particularly cholesterol released from astrocytes, is essential to determine the probability of the two main forms of synchronous vesicle release, UVR and MVR. Unexpectedly, we observed that MVR is >10 fold more sensitive to cholesterol levels than UVR, which provides an effective mechanism to modulate the MVR/UVR balance. Astrocytic cholesterol also modulates the spatial distribution of release events across individual AZs. Furthermore, astrocytic cholesterol supply regulates exo-/endocytosis coupling by controlling utilization of the two main forms of single-vesicle endocytosis (ultra-fast and fast) thus modulating the efficiency of vesicle uptake. These results suggest that via the release of cholesterol, astrocytes modulate and fine-tune the key spatiotemporal aspects of synaptic exo- and endocytosis in hippocampal excitatory synapses. Astrocytic supply of cholesterol thus adds a major regulatory pathway to the plethora of mechanisms by which astrocytes modulate synaptic transmission, and may present a new avenue towards alleviating synaptic deficits associated with pathological conditions of cholesterol imbalance.

### The role of astrocytic cholesterol in synaptic transmission

Astrocyte-derived cholesterol is widely known to be essential for synapse development and maintenance (42). During development, upregulation of genes involved in cholesterol biosynthesis in astrocytes precedes the activation of synaptic genes in neurons (43). Disruptions in cholesterol trafficking from astrocytes to neurons are associated with reduced number of excitatory synapses (44), and reduction in cholesterol impairs frequency of spontaneous synaptic activity in developing striatal neurons, consistent with reduction in synaptic connectivity (45). Despite this well-established role of astrocytic cholesterol in synapse development and maintenance, much less is known about its requirements for the efficiency of synaptic transmission. In particular, whether and to what extent astrocytic cholesterol is required for the different forms of vesicle release and/or uptake remains largely unknown. Our results show that astrocyte-produced cholesterol is critical for several key aspects of neurotransmitter release by determining the balance and spatial organization of different forms of vesicle exo- and endocytosis in hippocampal synapses. This form of modulation expands the known forms of astrocytic regulation mediated by release of gliotransmitters such as ATP, glutamate, or GABA, which occurs via exocytosis or release via variety of transporters and channels (46, 47). Gliotransmitters have been shown to up or down regulate efficiency of synaptic transmission via both pre- and postsynaptic modulation on timescales of tens of seconds to minutes (48-55). In addition, we recently reported a rapid astrocyte-dependent feed-back modulation of synaptic transmission mediated by activation of astrocytic mGluR5 and subsequent ATP release from astrocytes evoked by individual synaptic glutamate release events (15). ATP is rapidly degraded to adenosine, which activates synaptic A2A receptors to transiently potentiate MVR for a period of a few seconds. While the cholesterol-dependent changes in MVR/UVR balance occur on much slower time scales, the two forms of modulation could be complementary since cholesterol has been shown to regulate fusion pore conductance in astrocytes (56) and thus may regulate release of gliotrasmitters, such as ATP, thus effectively modulating MVR/UVR balance on multiple timescales. It remains to be determined what signaling pathways control astrocytic cholesterol release and whether changes in astrocytic cholesterol supply are triggered by neuronal activity.

It will also be important to determine in future studies the timescales on which changes in astrocytic cholesterol synthesis and release occur under normal and pathological conditions. Indeed, cholesterol imbalance has received a wide attention due to its hypothesized role in many neurodegenerative diseases (57). The function of astrocyte-derived cholesterol in modulating vesicle exo- and endocytosis may thus be relevant to pathophysiological conditions. Many disease states result in decreased brain cholesterol synthesis, for example in diabetes this occurs due to a reduction in sterol regulatory element-binding protein 2 (SREBP2)-regulated transcription (58). Altered brain cholesterol synthesis may thus contribute to the interaction between metabolic diseases, such as diabetes and altered brain function (58-60). Given our observations that astrocytic release of cholesterol is a potent modulator of synchronous forms of release and thus may regulate long-term changes in synaptic strength it will be important to examine in future studies to what extent changes in astrocytic cholesterol may contribute to pathological conditions associated with cholesterol imbalance.

### Cholesterol and the balance of different forms of vesicle release

Numerous previous studies have established that PM cholesterol depletion impairs synchronous synaptic transmission in a wide range of neurons (17, 19, 27, 35, 61-63). This reduction in synchronous release is often accompanied by increase in a spontaneous form of release (17-20) suggesting that lower cholesterol promotes spontaneous release at the expense of synchronous release. Notably, these previous observations were made at a whole-synapse level, and capacitance measurements of whole-synapse release in the Calyx of Held showed that cholesterol extraction reduces the magnitude of synaptic release predominately by decreasing the readily releasable pool (RRP) and the vesicle replenishment after RRP depletion (27). Consistent with this, we also observed that cholesterol extraction caused much larger reduction of the whole-synapse response evoked by high-frequency stimulation than the changes in release probability measured at the level of individual release events. Moreover, we observed a highly distinct sensitivity to changes in cholesterol of the two major forms of synchronous release. Specifically, MVR was markedly reduced by blocking cholesterol transport from astrocytes or its uptake by neurons, and it is nearly abolished in neurons grown in the absence of astrocytes, while UVR (and asynchronous release) are only mildly affected. Moreover, we found that supplementation of synapses with external cholesterol strongly enhanced MVR, while having no or little effects on UVR (or asynchronous release). Thus both up and down regulation of cholesterol are highly effective in modulating the UVR/MVR balance primarily by altering the occurrence of MVR.

Why are the two forms or synchronous release have such a different sensitivity to cholesterol levels? Mechanistically, cholesterol has been suggested to affect vesicle fusion via several mechanisms, most notably via syt1-induced membrane bending, by stabilizing fusion pores, or clustering of SNAREs or calcium channels (5). Cholesterol has also been implicated in reducing repulsive forces between membranes, thereby lowering the energy required for membrane fusion and fission processes (4), or by facilitating membrane stress relaxation involved in membrane bending (64). In ribbon synapses, cholesterol depletion impairs coupling between calcium channel opening and vesicle release by allowing calcium channels to move further from release sites (63). However, since all of these functions of cholesterol apply to both forms of release, the differential sensitivity to cholesterol is more likely to involve a specific aspect of vesicle fusion process that distinguishes MVR from UVR. One such distinction may arise from the differential spatial properties of the two forms of release. We previously found that UVR and MVR utilize a partially distinct subset of release sites, which are more specific to each form of release at the AZ periphery but more likely to overlap near the AZ center. Notably, clusters of presynaptic release machinery are smaller and less stable at the AZ periphery (65-67). Altering cholesterol levels could have stronger effects in smaller peripheral clusters and thus affect MVR to a larger extent. Future studies will be needed to elucidate the specific function of cholesterol that differentially affect the two forms of synchronous release.

### Cholesterol and the balance of different components of endocytosis

Cholesterol has long been known to be essential for clathrin-mediated endocytosis in a variety of non-neuronal cell types because clathrin is unable to induce curvature in the membrane depleted of cholesterol (68, 69). However, at the synapses, the specific role of cholesterol in endocytosis has been more elusive because of somewhat conflicting observations and the presence of several distinct and incompletely understood forms of endocytic mechanisms. Depletion of cholesterol has been shown to slow down uptake of FM1-43 in hippocampal terminals (61) and to reduce the rates of two forms of endocytosis, slow and fast, in the Calyx of Held terminals (27), but to increase spontaneous vesicle endocytosis in hippocampal synapses (17). Cholesterol extraction was also shown to slow down the overall kinetics of endocytosis after high-frequency stimulation in cortical cultured neurons, but cholesterol supplementation had no effect (28). Our whole-synapse measurements similarly show that cholesterol extraction slows the overall rate of endocytosis following high-frequency trains, while cholesterol supplementation had no measurable effect. However, it is important to consider major differences in the forms of endocytosis that are primarily utilized in response to strong high-frequency stimulation and those coupled with individual vesicle release events. In our measurements, single-vesicle release events are coupled to either ultra-fast (∼100 ms) or fast (∼500ms-1 s) forms of endocytosis, while slow endocytosis (tens of seconds) is not detectable, but could occur in no more than ∼3% of all events in which no measurable decay is detected over 1 s period. In contrast, slow and fast forms of endocytosis are predominately observed after high-frequency trains in whole-synapse measurements, while ultra-fast endocytosis is not utilized or detectable. In line with these differences, our single-vesicle measurements demonstrate highly distinct roles of cholesterol in regulating the two main forms of endocytosis coupled with individual release events. We observed that either extraction of cholesterol, or inhibition of the release/uptake of astrocytic cholesterol increases the occurrence of ultra-fast endocytosis while reducing the occurrence of fast endocytosis, and the opposite is observed with cholesterol supplementation. Via these opposing changes, cholesterol levels thus fine-tune the balance and preferential utilization of ultra-fast vs fast forms of endocytosis coupled with single release events.

The mechanisms underlying fast and particularly ultra-fast endocytosis are not well understood, making it challenging to pinpoint the origins of the differential effects of cholesterol on these two forms of endocytosis. Recent studies suggested the existence of pre-formed omega pits as a possible mechanism of ultra-fast endocytosis (70). The limited number of such pre-formed pits could explain why this fastest and energetically more favorable form of endocytosis is only utilized in ∼20-25% of release events. Recent studies also suggested a possibility that ultrafast endocytosis is caused by a mechanical membrane displacement that aids rapid vesicle formation at the AZ peripheral junctions with actin cytoskeleton (71). However, this mechanism requires release of two or more vesicles i.e. MVR, while the opposing sensitivity to cholesterol of the two forms of endocytosis was observed for all release events. We previously found several major mechanistic differences between the two forms of endocytosis, including their calcium-dependence and requirements for dynamin (29), but whether these differences may contribute to the effects of cholesterol is not clear. Deciphering the mechanisms by which cholesterol differentially affects the two forms of endocytosis is further complicated by the complex relationship between cholesterol and membrane curvature (3, 69). During spontaneous curvature, cholesterol behaves as an extremely negative curvature lipid in disordered membranes but appears to have a net positive effect in the presence of saturated lipids (3, 69), so its effects may also depend on local membrane lipid composition. While the precise molecular mechanisms governing the opposing effects of cholesterol on the two forms of endocytosis will require future investigation, our results uncover a critical role of astrocytic cholesterol in modulating the exo- /endocytosis coupling of individual release events in central synapses. This fine-tuning of the balance between different forms of exo- and endocytosis may represent a previously unrecognized form of astrocyte-dependent plasticity that maybe relevant to disorders of brain cholesterol imbalance.

## ASKNOLEDGMENTS

This work was supported by an NIH grant R35 NS111596 to VAK.

## MATERIALS AND METHODS

### KEY RESOURCE TABLE

| REAGENT or RESOURCE | SOURCE | IDENTIFIER |
| --- | --- | --- |
| **Bacterial and Virus Strains** |  |  |
| pFU-vGluT1-pHGFP-W | Viral Vectors Core at Washington University | N/A |
| **Biological Samples** |  |  |
| Hippocampus of Long-Evans rat pups | Charles River | Cat. No. 006 |
| **Chemicals, Peptides, and Recombinant Proteins** |  |  |
| APV | Sigma-Aldrich | Cat.No. 79055-68-8 |
| CNQX disodium salt hydrate | Sigma-Aldrich | Cat.No. 115066-143 |
| HEPES | Sigma-Aldrich | Cat.No. 7365-45-9 |
| D-(+)-Glucose | Sigma-Aldrich | Cat.No. 50-99-7 |
| Calcium chloride dihydrate | Sigma-Aldrich | Cat.No. 10035-04-8 |
| Magnesium chloride hexahydrate | Sigma-Aldrich | Cat.No. 7791-18-6 |
| Minimum Essential Media (MEM) -No phenol Red | Gibco™ | Cat.No. 51200038 |
| Characterized Fetal Bovine Serum | Gibco™ | Cat.No. 10437028 |
| Penicillin-Streptomycin (5, 000 U/ml) | Gibco™ | Cat.No. 15070063 |
| N-21 | R&D systems | Cat No. AR008 |
| FBS | Gibco | Cat No. 26140-079 |
| N-2 Supplement (100x) | Gibco™ | Cat.No. 17502048 |
| Donor Equine Serum | HyClone | Cat.No. SH30074.03 |
| Sodium Pyruvate (100 mM) | CORNING | Cat.No. 25-000-CI |
| Neurobasal-A Medium | Gibco™ | Cat.No. 12349015 |
| B-27 supplement (50x), serum free | Gibco™ | Cat.No. 17504044 |
| GlutaMAX Supplement | Gibco™ | Cat.No. 35050061 |
| Sodium pyruvate | Sigma-Aldrich | Cat.No. P2256 |
| Collagen Type I, Rat Tail | CORNING | Cat.No. 354236 |
| Cover Glasses | Thermo Fisher Scientific | Cat.No. 12-545-80 |
| Papain | Worthington Biochemical | Cat.No. LS003126 |
| PDL(poly-D-lysine) | BD Biosciences | Cat.No. 40210 |
| Trypsin-EDTA (0.05%), phenol red | Gibco™ | Cat.No. 25300054 |
| **Lovastatin** | Sigma-Aldrich | Cat.No. 75330-75-5 |
| **Probucol** | Sigma-Aldrich | Cat.No. 23288-49-5 |
| **Valspodar** | Sigma-Aldrich | Cat.No. 121584-18-7 |
| **PCSK9** | abcam | Cat.No. ab167759 |
| **U18666A** | Sigma-Aldrich | Cat.No. 3039-71-2 |
| **LDL** | Thermo Fisher Scientific | Cat.No. L3486 |
| **MβCD** | Sigma-Aldrich | Cat.No. 128446-36-6 |
| **WSC** | Sigma-Aldrich | Cat.No. C4951 |
| **Recombinant DNA** |  |  |
| **vGlut1-pHluorin** | (25) | N/A |
| **pGP-CMV-jGCaMP8f** | Addgene | 162373 |
| **Software and Algorithms** |  |  |
| **MATLAB** | MathWorks | RRID: SCR_001622 |
| **u-track 2.0** | https://www.utsouthwestern.edu/labs/danuser/software/#utrack_anc | Gaudenz Danuser Lab |
| **ImageJ** | https://imagej.nih.gov/ij/ | RRID: SCR_003070 |

### RESOURCE AVAILABILITY

#### Lead contact

Further information and requests for resources and reagents should be directed to and will be fulfilled by the Lead Contact, Dr. Vitaly A. Klyachko.

#### Materials availability

This study did not generate new or unique reagents or other materials.

#### Data and code availability

- This paper does not report standardized data types. All data reported in this paper will be shared by the lead contact upon request.
- This paper does not report standalone custom code. MATLAB was used to appropriately organize, process, and analyze data and corresponding routines are available from the lead contacts upon request.
- Any additional information required to reanalyze the data reported in this paper is available from the lead contacts upon request.

### EXPERIMENTAL MODEL AND SUBJECT DETAILS

#### Animals

All animal process conformed to the guidelines approved by the Washington University Animal Studies Committee. Mixed sex embryos from wild-type rats of the Sprague-Dawley strain were used for primary neuronal cultures.

#### Neuronal Cell Culture

Neuron culture were performed with hippocampus of rat of mixed sexes as previous described (15, 29, 30). Briefly, hippocampi were dissected from E16-17 pups, dissociated by papain digestion, and plated on poly-d-lysin coated glass coverslips. Neuron cultured in Neurobasal media supplemented with B-27 supplement. Neurons were grown on top of a confluent astrocyte monolayer. For neuron-only culture, dissociated cells were treated with CultureOne Supplement to remove proliferating astrocytes as described (15).

### METHOD DETIALS

#### Lentiviral Infection

vGlut1-pHluorin were generously provided by Drs. Robert Edwards and Susan Voglmaier (UCSF) (25). Lentiviral vectors were generated by the Viral Vectors Core at Washington University. SF-iGluSnFR(A184) were kindly made available by Dr. Loren Looger (Addgene viral Prep #106174-AAV1) (34). Lentiviral infection was performed on hippocampal neuron culture at DVI3 as described (15, 29, 30).

#### Near-TIRF microscopy

All near-total internal reflection fluorescence (near-TIRF) experiments were performed at 37° within a whole-microscope incubator chamber (OTKAI HIT) as described (15, 29, 30). vGltu1-pHluorin was excited by a 488 nm laser (Cell CMR-LAS-488, Olympus). Images were captured using an inverted microscope (IX83, Olympus) and a 150x/1.45NA objective lens. The nano-precision TIRF-based Z-drift compensation system (IX-ZDC) was used to keep constant position of the focal plan during imaging process. Near-TIRF with penetration depth of <1 µm was achieved by adjusting the incident angle to 63.7°, which is near the critical angle of 63.6°. Images were captured every 50 ms (with exposure time of 49.38 ms) with a cooled EMCCD camera (iXon life 888, ANDOR). Field stimulation was performed by using a pair of platinum electrodes and controlled by software via Master-9 stimulus generator (A.M.P.I.). Samples were perfused with bath solution containing 125 mM NaCl, 2.5 mM KCl, 2 mM CaCl_2_, 1 mM MgCl_2_, 10 mM HEPES, 30 mM Glucose, 50 µM D,L-2-amino-5phosphonovaleric acid (APV), 10 µM CNQX, adjusted to pH 7.4.

#### Pharmacology

Lovastatin, probucol, valspodar, and PCSK9 were diluted in dimethyl sulfoxide (DMSO) and stored at -20°C. U18666A was diluted in DMSO and stored at 4°C. Samples were pre-incubated with 10 µM lovastatin, 10 µM probucol, 5 µM valspodar, 50 nM PCSK9, or 1 µM U18666A for 2 days for prior to the beginning of the recording. Control conditions were treated with same final concentration of DMSO. MβCD and WSC solutions were prepared on the day of imaging and applied for 10 min before imaging.

### QUANTIFICATION AND STATISTICAL ANALYSES

#### Event detection and localization

vGlut1-pHluorin-based release event detection and localization at subpixel resolution were achieved as described previously (15, 29, 30) using MATLAB and uTrack software package, which was kindly made available by Dr.Gaudenz Danuser’s lab (72, 73). Localization precision was determined directly from least-squares Gaussian fits of individual events as described (15, 29, 30). To determine the general Pr, boutons were stimulated at 1 Hz and Pr was determined by calculating the probability of detecting release events as determined over a 200 s period.

#### Detection of MVR

MVR events were identified as we described previously (15, 30) by calculating the mean and standard deviation values of event peak intensities determined individually for each synapse during 1 Hz, 200 s stimulation period. Events with transient intensities exceeding the sum of the mean and twice the standard deviation were designed as MVR, while the remaining events were classified as UVR.

#### Definition of AZ release sites/clusters

Release sites/clusters were defined using hierarchical clustering algorithm with a clustering diameter of 50 nm using built-in functions in MATLAB as described (15, 29, 30). We have previously shown that the observed cluster do not arise from random distribution of release events but rather represent a set of defined and repeatedly reused clusters within the AZs (38).

#### Analysis of endocytosis

The ratio of ultra-fast/fast endocytosis, which we refer to as ultrafast ratio, was defined by comparing the areas of the Gaussian fits to endocytosis tau histogram of fast vs. ultra-fast components as we described previously (30). Briefly, bi-Gaussian curves were used to fit the distribution of decay tau and the areas under each of the two Gaussian curves corresponding to the ultra-fast and fast endocytosis events were determined. The ultrafast ratio was defined by dividing the area corresponding to the ultrafast component by the sum of ultrafast and fast areas.

To determine the fraction of retrieval, the decay of individual event traces was fitted using a mono-exponential curve. The fraction of retrieval was determined by dividing the difference of the amplitude values of the curve at peak and at the end of the 1s period following the stimulation by the event’s peak amplitude, as described (30). Calculated fractions of retrieval were categorized into four groups: “partial” and “excessive” retrieval corresponded to fraction <0.8 or >1.2, respectively. Fraction between 0.8 and 1.2 were categorized as “quantal”, and “no retrieval” was determined if no detectable signal decay was observed during the 1 s period.

The dwell time was determined from a fit to the individual event traces using a combination of plateau and exponential decay. Dwell time and half-time were calculated by determining the plateau periods before the decay and time to reach half of the peak values, respectively.

#### Data inclusion and exclusion criteria

A minimum of 10 detected release events per bouton was required of all spatial and temporal analyses of exo- and endocytosis to ensure an adequate determination of AZ dimensions and adequate sampling of event spatiotemporal features across individual AZs during 1 Hz stimulation for 200 s. For all experiments, including the determination of Pr, functional synapses were defined as those with a Pr ≥ 0.05; synapses with a Pr below this threshold were excluded from analysis.

#### Statistical analysis

Statistical analyses were performed in MATLAB. Statistical significance was determined using a two-tailed t test, Tukey-Kramer ANOVA, and Kolmogorov-Smirnov (K-S) tests where appropriate. Statistical tests used to measure significance, the corresponding significance level (*p*-value), and the values of n are provided for each panel in **Table S1**. Due to the very large number of data points per measurement, individual data points are not shown in plots; the mean ± SEM values are reported and plotted. *p* < 0.05 was considered statistically significant.

## SUPPLEMENTARY FIGURE LEGENDS

**Supplementary Figure 1.**
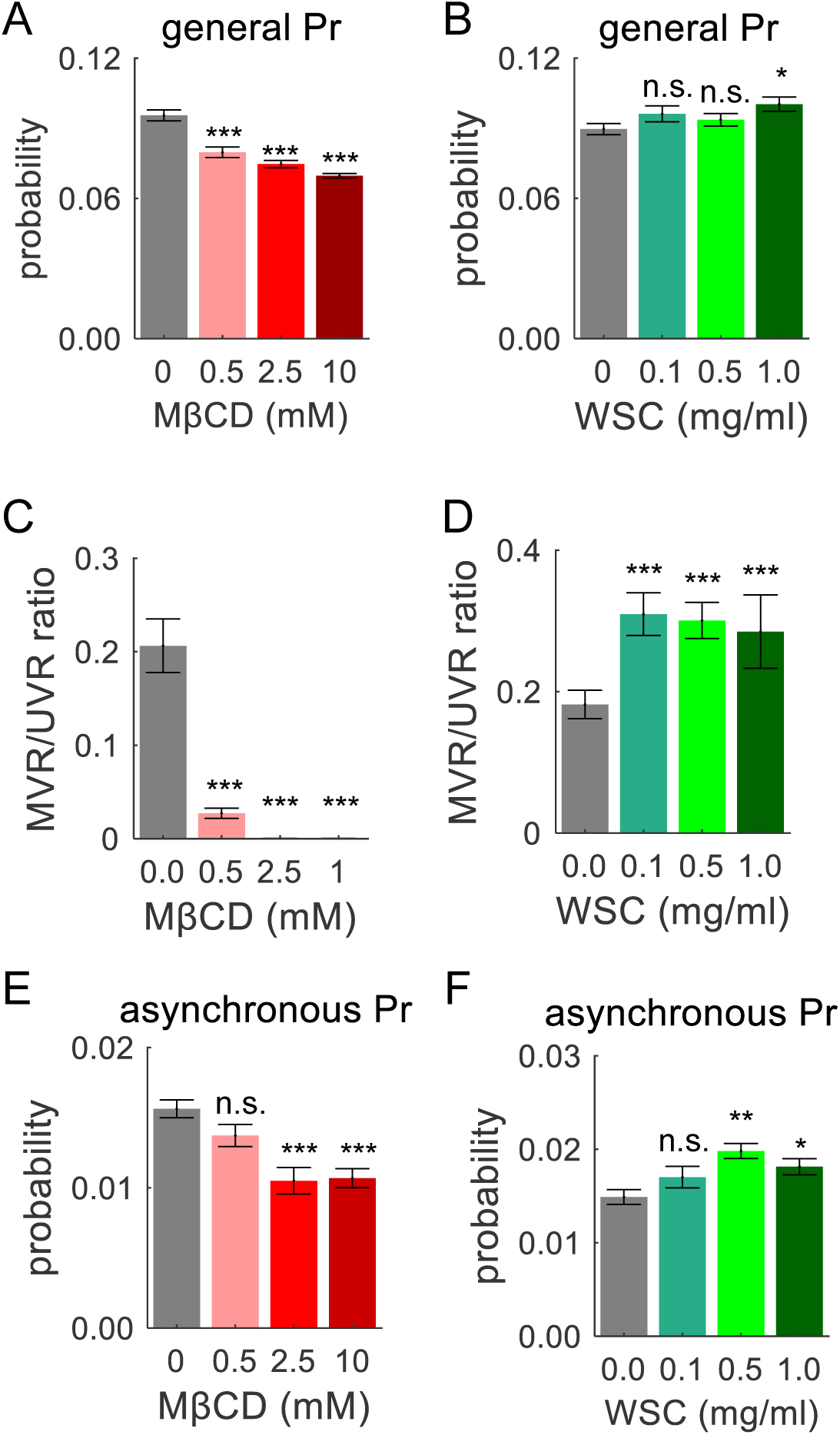
The effects of cholesterol on release probability and asynchronous release. **(A, B)** Overall Pr calculated for individual synapses stimulated at 1 Hz for 200 seconds under varying concentrations of MβCD (A) or WSC (B). **(C, D)** Average MVR/UVR ratio in individual synapses in varying concentrations of MβCD or WSC. **(E, F)** Pr of asynchronous vesicle in varying concentrations of MβCD or WSC.

**Supplementary Figure 2.**
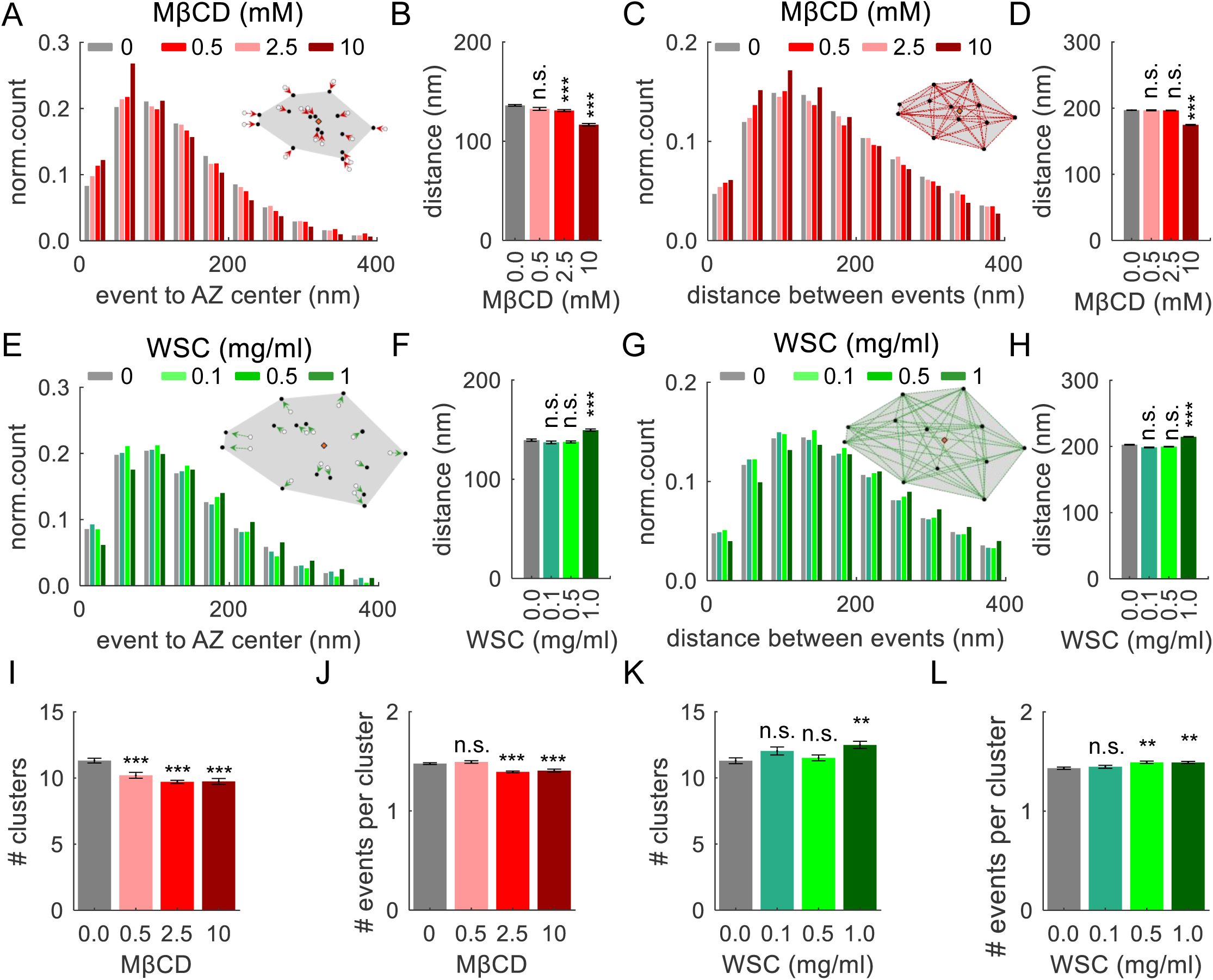
Cholesterol levels regulate spatial localization of vesicle release across the AZ. **(A)** Distribution of distances from release events to the AZ center within individual synapses under varying concentrations of MβCD. **(B)** Average distances of release events to the AZ center under varying concentrations of MβCD from data in (**A**). **(C)** Distributions of distances between release events within individual synapses under varying concentrations of MβCD. **(D)** Average distances between release events within individual synapses under varying concentrations of MβCD from data in (**C**). **(E-H)** Same analysis as (**A-D**), but with varying concentrations of WSC. **(I)** Average number of clusters/release sites detected in individual AZs under different concentrations of MβCD. **(J)** Average number of events detected per cluster under different concentrations of MβCD. **(K, L)** Same analysis as (**I-J**), but with varying concentrations of WSC.

**Supplementary Figure 3.**
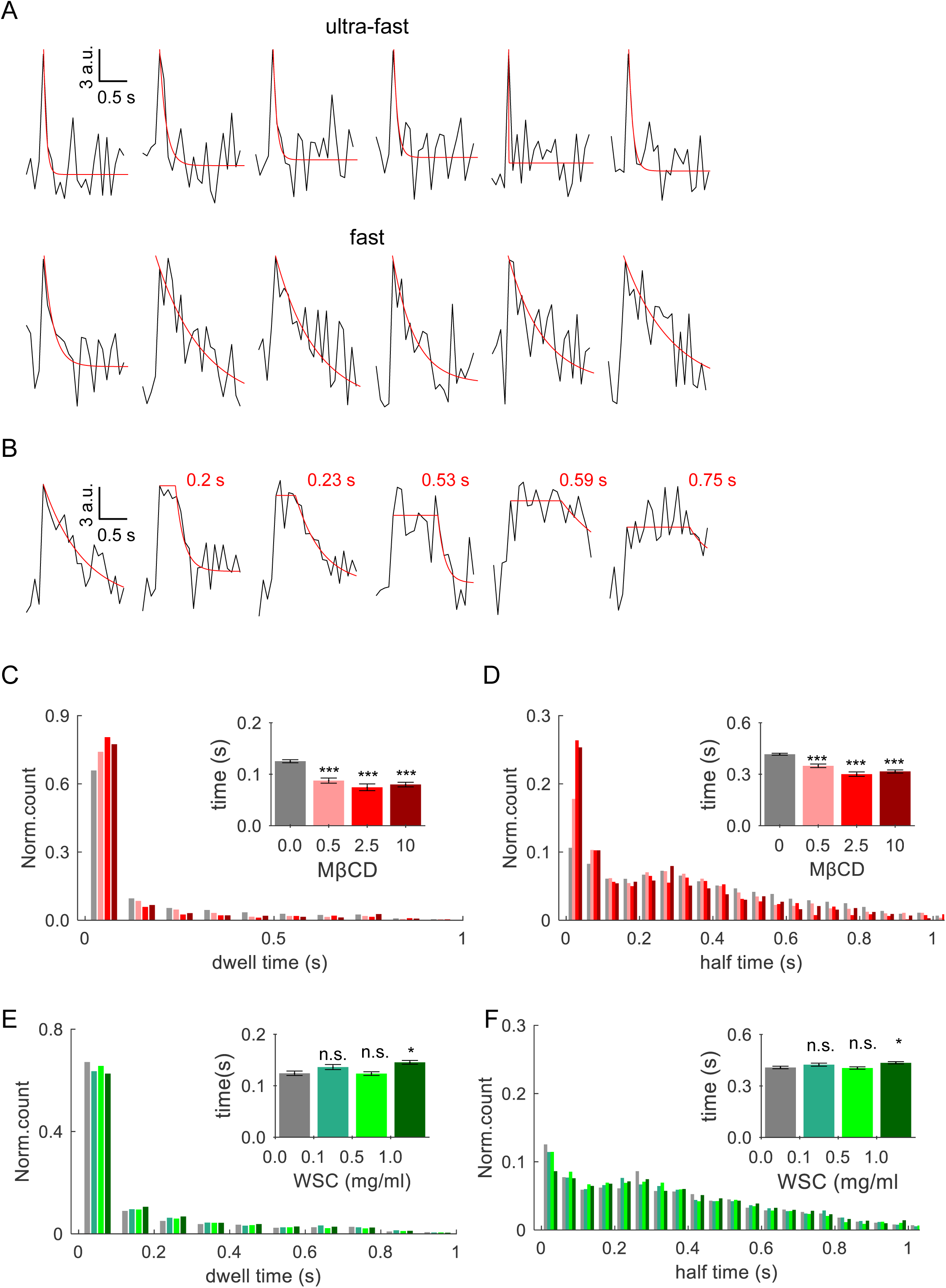
Cholesterol levels regulate preferential mode and dwell time of endocytosis. **(A)** Sample traces of single vesicle events evoked by 1-AP stimulation, illustrating two distinct rates of endocytosis: ultrafast (top row) or fast (bottom row). **(B)** Sample traces of single-vesicle events with varying endocytosis dwell time. **(C)** Distribution of dwell time of single-vesicle release events in different concentrations of MβCD. *Inset:* corresponding average dwell time in different concentrations of MβCD. **(D)** Same as (**C**) for the event decay half-time. **(E,F)** Same analyses as (**C, D**), but for varying concentrations of WSC.

**Supplementary Figure 4.**
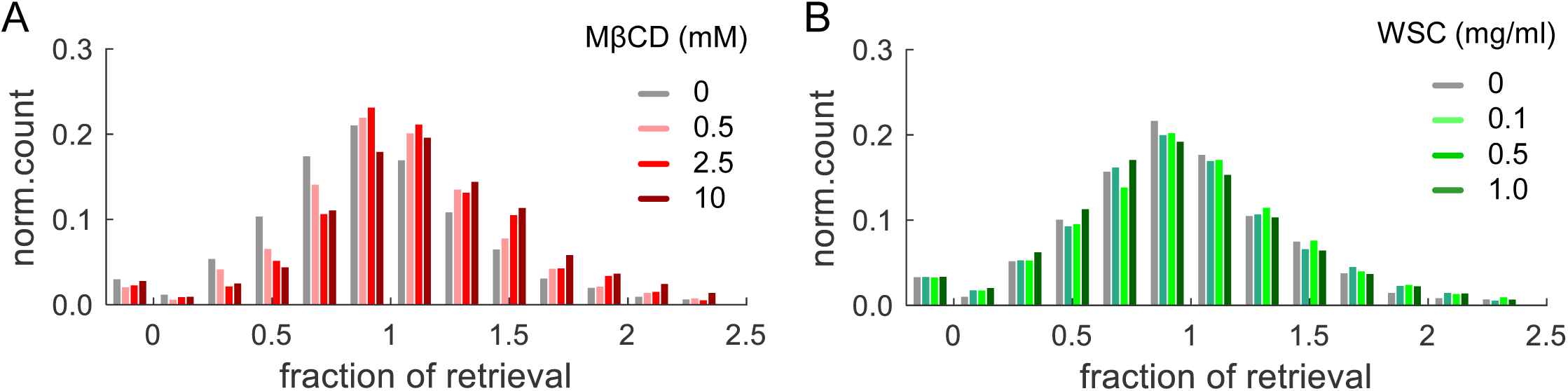
Cholesterol levels regulate the extent of membrane retrieval during endocytosis. (**A,B**) Distributions of the fraction of retrieval in different concentrations of MβCD (**A**) or WSC (**B**).

**Supplementary Figure 5.**
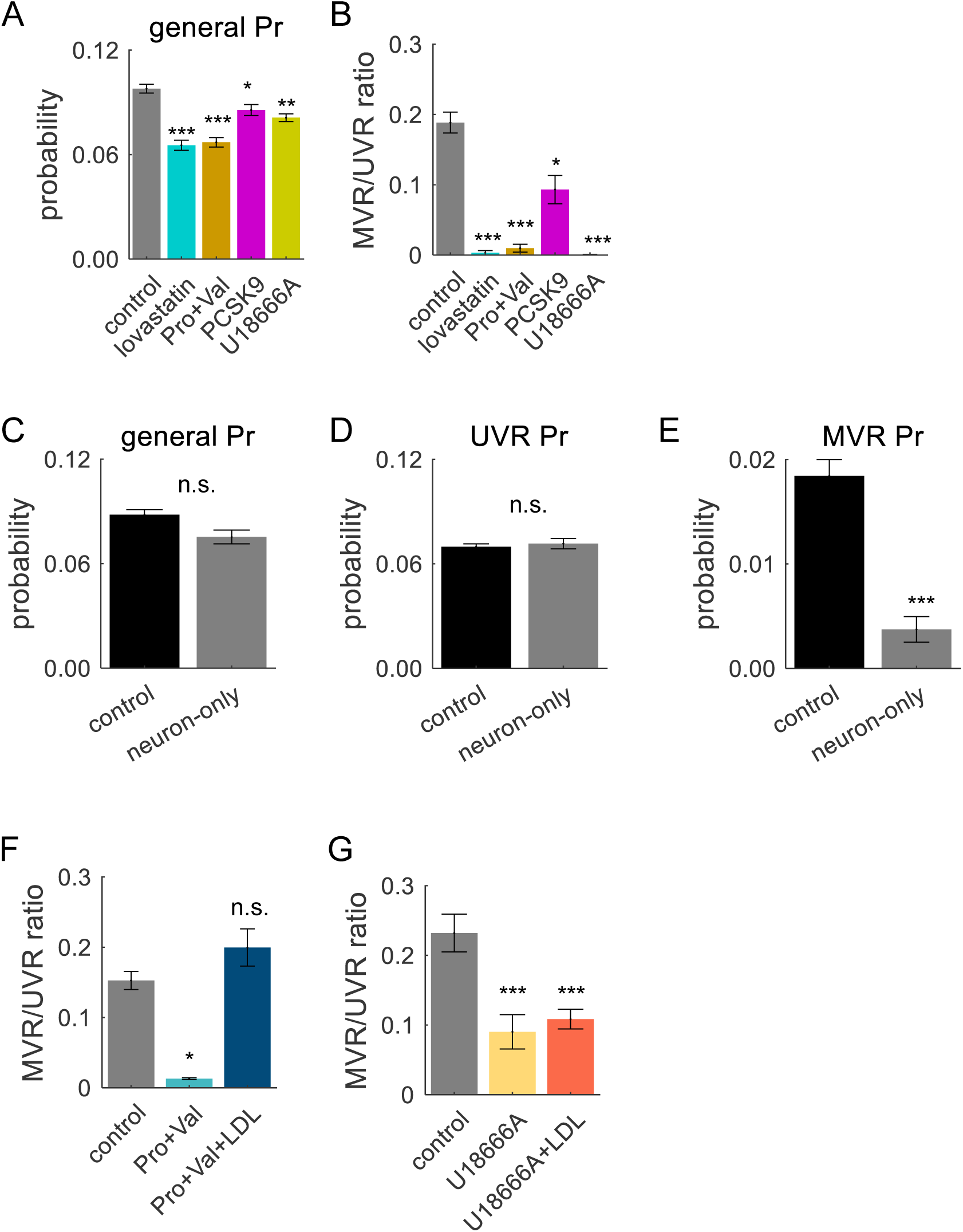
Astrocytic Cholesterol Determines Release Probability and MVR/UVR Ratio. **(A)** Pr in individual synapses, calculated based on detection of all release events evoked at 1 Hz for 200 seconds under various conditions blocking astrocytic cholesterol delivery pathway to neurons (same as in Fig 3), as indicated. **(B)** MVR/UVR ratio in individual synapses in the same conditions as in (**A**). **(C,D,E)** Pr overall, and Pr of UVR and MVR, in neurons cultured with (control) or without (neuron-only) an astrocyte layer. **(F)** MVR/UVR ratio in individual synapses pre-incubated with probucol and valspodar, with or without LDL. **(G)** Same as in (**F**), but for U18666A, with or without LDL.

**Supplementary Figure 6.**
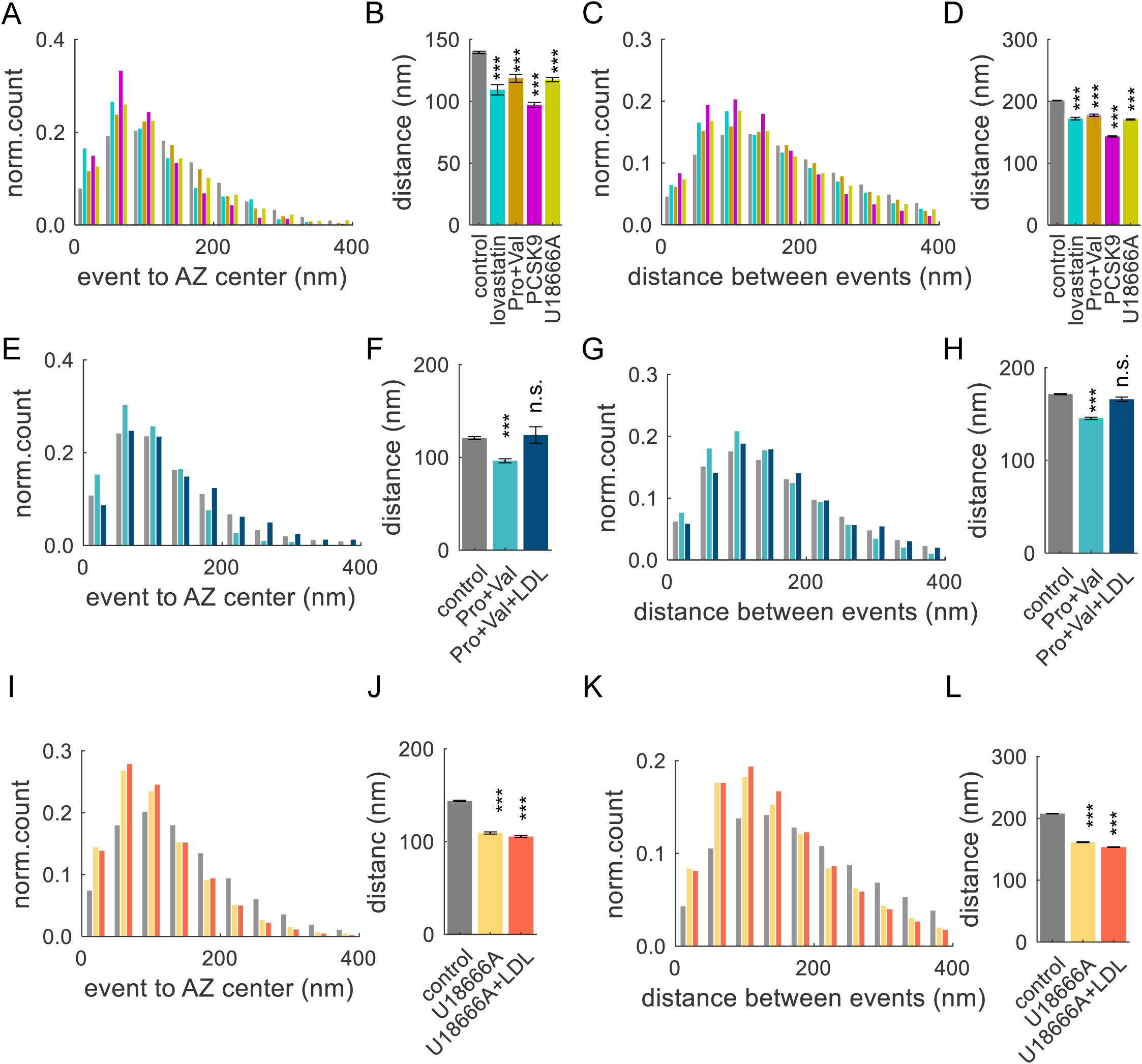
Astrocytic Cholesterol Determines Spatial Distribution of Vesicle Release Across the AZ. **(A)** Distribution of distances from release events to the AZ center recorded following 2-day pre-incubation with various inhibitors that disrupt the pathway of cholesterol delivery from astrocytes to neurons, as shown in **Fig 3A**. **(B)** Average distances of release events to the AZ center in data from (**A**). **(C)** Distributions of distances between release events within individuals synapses under the same conditions as in (**A**). **(D)** Average distances between release events within individual synapses in data from (**C**). **(E-H)** Same analyses as in (**A-D**), but with probucol and valspodar, with or without LDL. **(I-L)** Same analyses as in (**A-D**), but with U18666A, with or without LDL.

## SUPPLEMENTARY TABLE LEGEND

**Table S1.**
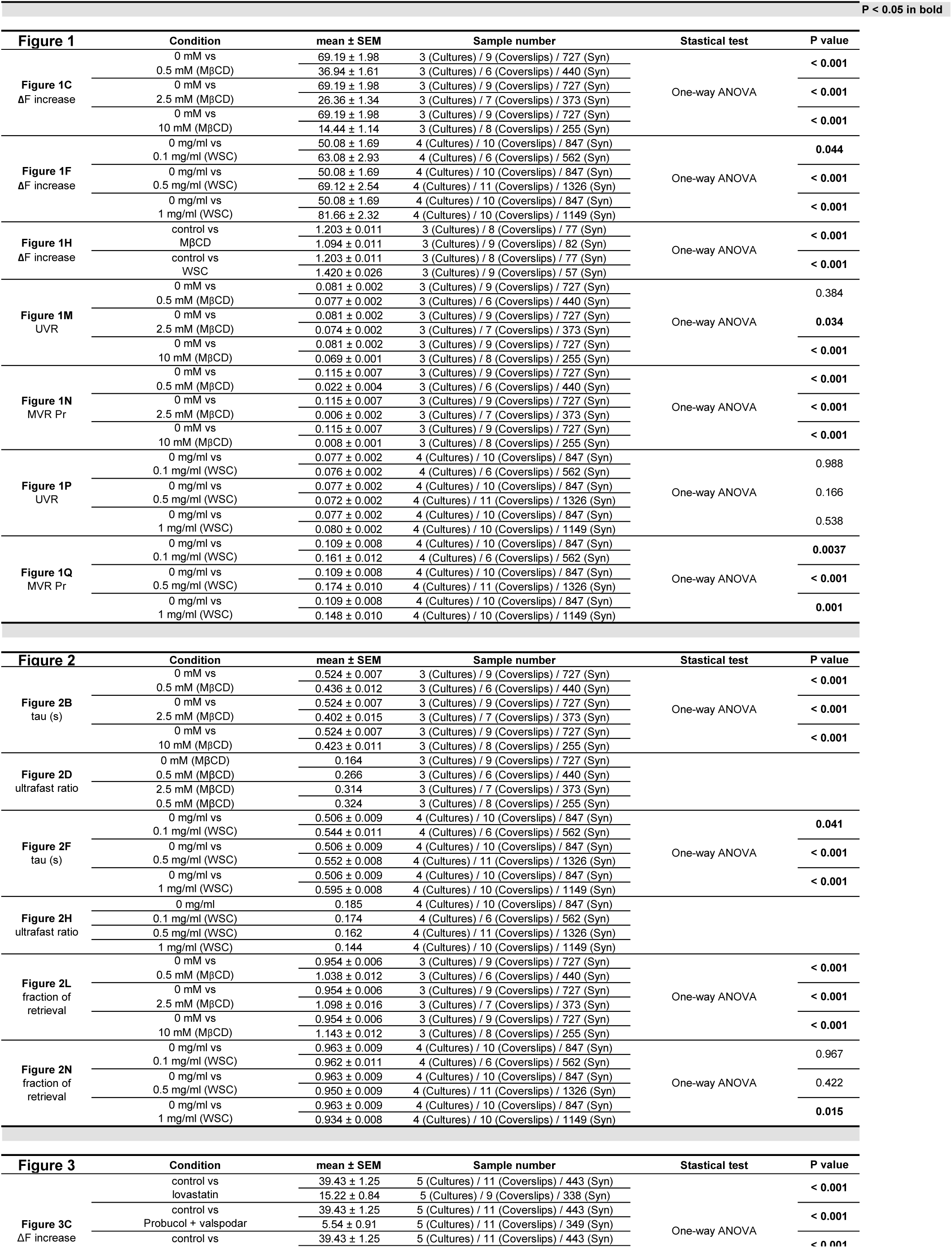
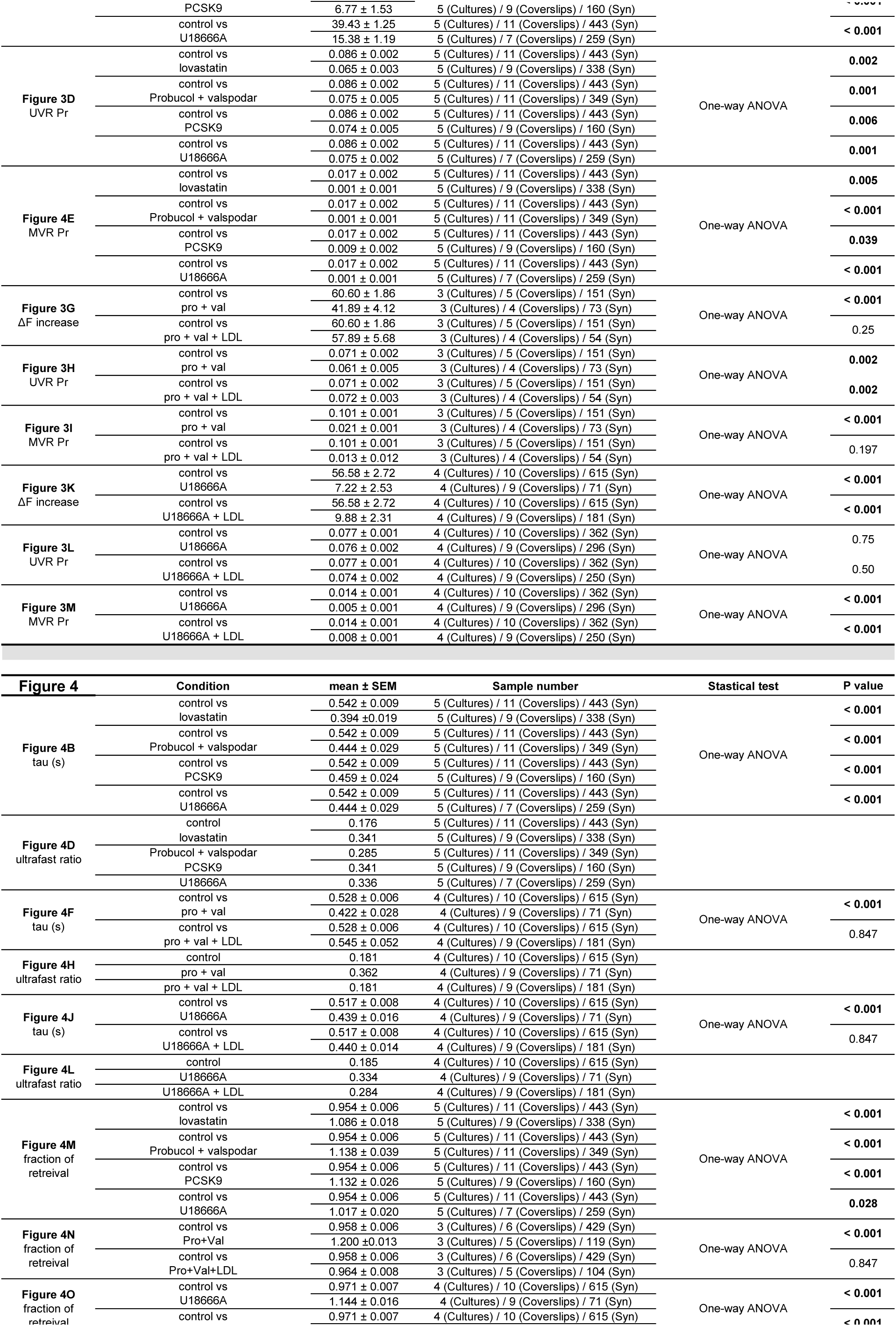
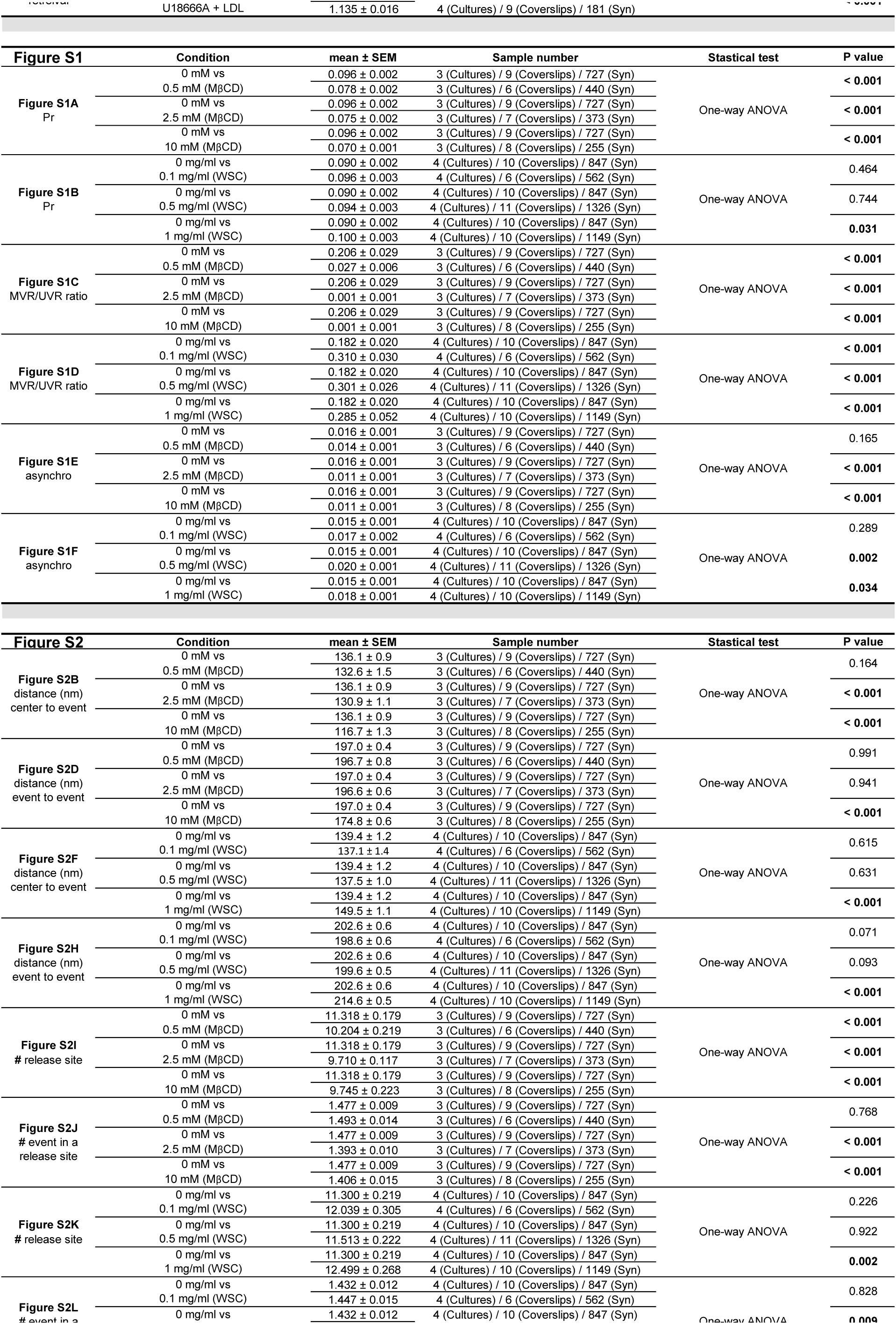
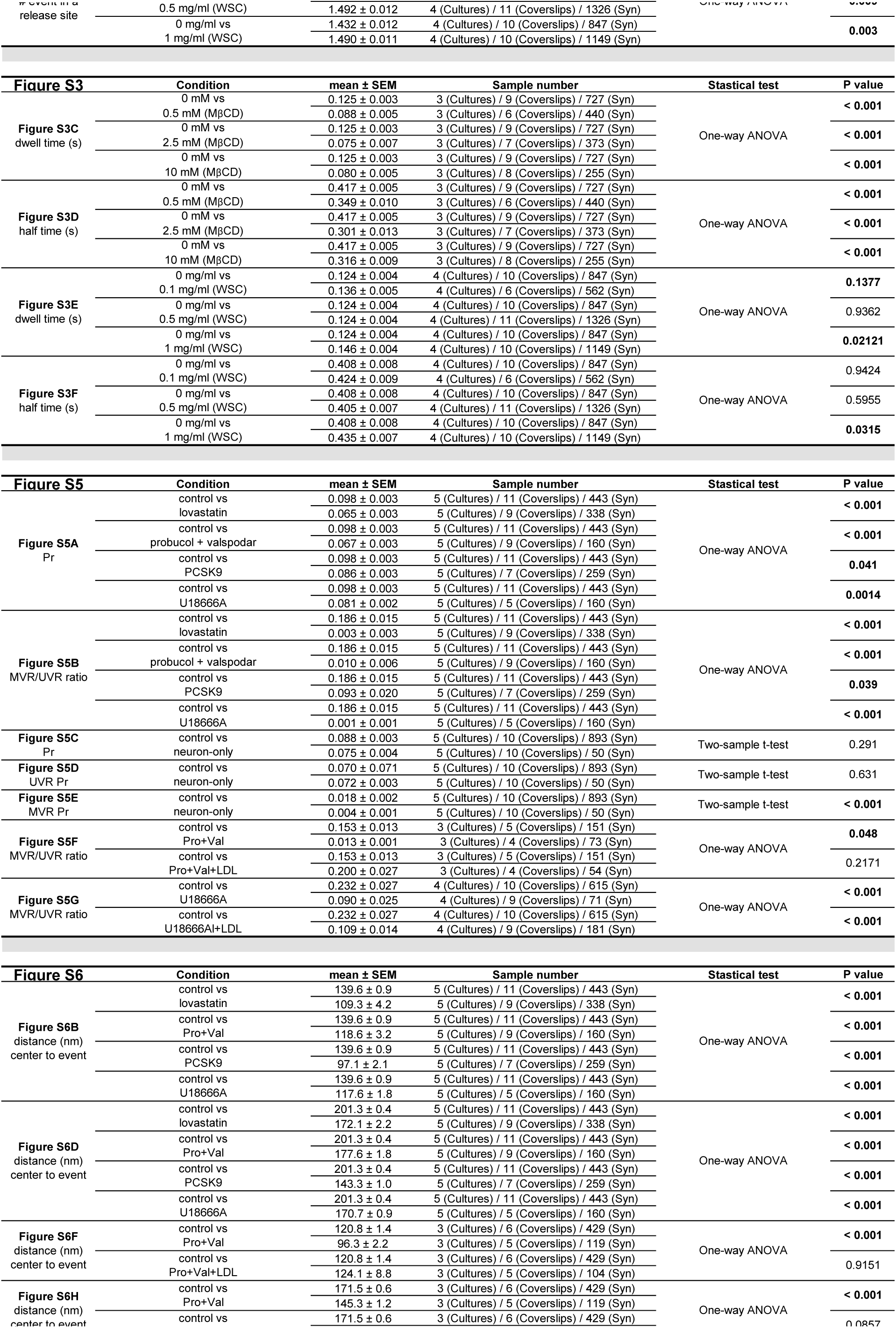
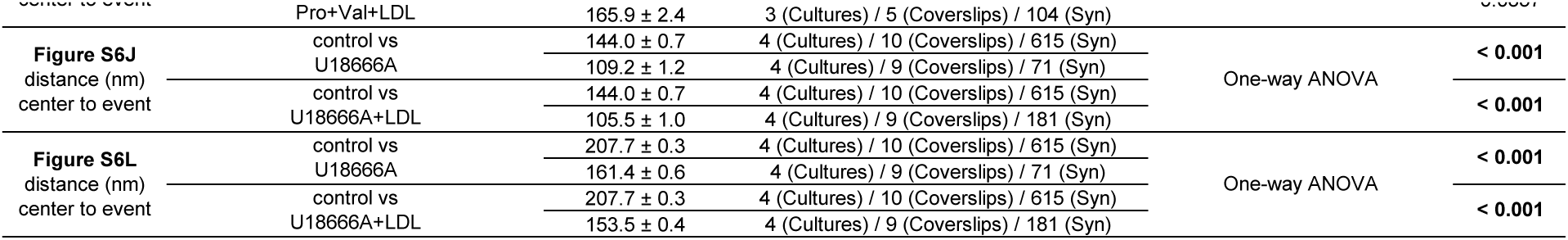
Statistical data for all figures. *Mean, SEM, number of experiments, coverslips and synapses, statistical test and P values are shown for every panel in each figure*.

